# Metabolomic profiles of stony coral species from the Dry Tortugas National Park display inter- and intraspecies variation

**DOI:** 10.1101/2024.10.08.617290

**Authors:** Jessica M. Deutsch, Alyssa M. Demko, Olakunle A. Jaiyesimi, Gabriel Foster, Adelaide Kindler, Kelly A. Pitts, Tessa Vekich, Gareth Williams, Brian K. Walker, Valerie J. Paul, Neha Garg

**Affiliations:** School of Chemistry and Biochemistry, Engineered Biosystems Building, Center for Microbial Dynamics and Infection, Georgia Institute of Technology, Atlanta, GA, USA; Smithsonian Marine Station, Smithsonian Institution, Fort Pierce, FL, USA; School of Ocean Sciences, Bangor University, Anglesey, UK; GIS and Spatial Ecology Laboratory, Halmos College of Arts and Sciences, Nova Southeastern University, Dania Beach, FL, United States

**Keywords:** Comparative metabolomics, Scleractinia, stony coral tissue loss disease, Symbiodiniaceae, tocopherol quinones, acylcarnitines

## Abstract

Coral reefs are experiencing unprecedented loss in coral cover due to increased incidence of disease and bleaching events. Thus, understanding mechanisms of disease susceptibility and resilience, which vary by species, is important. In this regard, untargeted metabolomics serves as an important hypothesis-building tool enabling delineation of molecular factors underlying disease susceptibility or resilience. In this study, we characterize metabolomes of four species of visually healthy stony corals, including *Meandrina meandrites*, *Orbicella faveolata*, *Colpophyllia natans*, and *Montastraea cavernosa*, collected at least a year before stony coral tissue loss disease reached the Dry Tortugas, Florida and demonstrate that both symbiont and host-derived biochemical pathways vary by species. Metabolomes of *Meandrina meandrites* displayed minimal intraspecies variability and highest biological activity against coral pathogens when compared to other species in this study. Application of advanced metabolite annotation methods enabled delineation of several pathways underlying interspecies variability. Specifically, endosymbiont-derived vitamin E family compounds, betaine lipids, and host-derived acylcarnitines were among the top predictors of interspecies variability. Since several metabolite features that contributed to inter- and intraspecies variation are synthesized by the endosymbiotic Symbiodiniaceae, which could be a major source of these compounds in corals, our data will guide further investigations into these Symbiodiniaceae-derived pathways.

**Importance.:** Previous research profiling gene expression, proteins, and metabolites produced during thermal stress has reported the importance of endosymbiont-derived pathways in coral bleaching resistance. However, our understanding of interspecies variation in these pathways among healthy corals and their role in diseases is limited. We surveyed the metabolomes of four species of healthy corals with differing susceptibilities to the devastating stony coral tissue loss disease and applied advanced annotation approaches in untargeted metabolomics to determine the interspecies variation in host and endosymbiont-derived pathways. Using this approach, we propose the survey of immune markers such as vitamin E family compounds, acylcarnitines, and other metabolites to infer their role in resilience to coral diseases. As time-resolved multi-omics datasets are generated for disease-impacted corals, our approach and findings will be valuable in providing insight into the mechanisms of disease resistance.

## Introduction

Coral reefs are an essential component of marine ecosystems, providing habitats for nearly a quarter of ocean life(1, 2), prevent shoreline erosion, and contribute to local economies(3, 4) and cultural practices(5). Climate change and increases of anthropogenic stressors have resulted in an unprecedented increase in incidence of coral bleaching and exposure to diseases(6–9). Diseases may result in coral colony mortality and can drive local extinction of corals(10). One of the biggest threats to Caribbean reefs is the continued spread of stony coral tissue loss disease (SCTLD). This virulent disease spread rapidly across Florida’s Coral Reef over the course of several years, affecting ∼22 scleractinian coral species since its first observation in 2014(11), and has since spread throughout the Caribbean(12). Disease susceptibility, resilience, and lethality vary significantly among affected species(11, 13–16), but underlying mechanisms behind resilience remain unknown.

The Scleractinian (stony coral) coral holobiont is diverse, consisting of the coral animal (host), the endosymbiotic dinoflagellate algae, archaea, bacteria, viruses, and fungi(17–20). Many coral species are sessile organisms that rely heavily on symbiosis with photosynthetic microalgae (Symbiodiniaceae) for their daily energy requirement and through associations with microbial communities for nutrition and defense(17, 21–25). Within the holobiont, multiple interactions occur among all members. The coral animal provides a physical habitat for the endosymbiont and a diversity of microorganisms within the holobiont. Nutrient exchange between the coral host and Symbiodiniaceae include the exchange of phosphorus and nitrogen (from the coral animal), oxygen, and carbon (from the Symbiodiniaceae, generated through photosynthesis). Studies have shown that Symbiodiniaceae can modulate intra- and interspecies susceptibility to bleaching(26, 27) and disease(28). Metabolite exchange between endosymbiotic zooxanthellae and bacteria influence the fitness of the endosymbiont(29). Bacteria selected by the coral holobiont fulfill several key roles in holobiont metabolite cycling and pathogen defense(30). *Endozoicomonas*, for example, are proposed to provide vitamins to both endosymbiotic dinoflagellates and the coral host(24). The bacterial communities are uniquely structured between the coral skeleton, tissue, and mucus(25). While the microbial community composition within the skeleton and tissue are associated with the coral host species, environmental factors may have a greater impact on the community structure within coral mucus(25). Beneficial bacteria (symbionts) also produce natural products that can provide competitive advantages to the coral holobiont and aid in responding to pathogens(31). Studies that profiled heat-sensitive and heat-tolerant corals have established that symbiosis as well as heterotrophy play a key role in determining which species thrive in the face of increasing ocean temperatures and that endosymbiont identity is an important factor in survival(32–37). Although the cause of SCTLD remains unknown, dysbiosis in the coral holobiont occurs with breakdown in the host-endosymbiont relationship in the gastrodermis resulting in necrosis and opportunistic infections by bacterial pathogens(38). There is also evidence of in situ symbiophagy (symbiont degradation) by the coral host in SCTLD-exposed corals(39, 40).

‘Omics techniques such as transcriptomics, proteomics, and metabolomics hold promise for delivering insights into biochemical pathways that may drive differences in disease response(41). Pairing multiple ‘omics techniques enabled key insights into how *Endozoicomonas* can provide key immune response related metabolites (such as vitamins) to the coral host and symbiotic zooxanthellae within the coral holobiont(24). Additionally, by comparing transcriptomes of stony corals of species *Acropora hyacinthus* either resilient or sensitive to bleaching stress, expression of genes important for survival even under non-stress conditions in a phenomenon termed frontloading was observed(42). The differential expression of orthologs related to vesicular trafficking and signal transduction were positively correlated to species-specific susceptibility to SCTLD(39).With advancements in data annotation strategies, untargeted metabolomics is also being increasingly employed to generate, refine, and validate hypotheses to untangle interactions between different members of the holobiont(43, 44). Metabolomics has been largely applied to profile different genotypes of corals(45), locations(46, 47), delineate biochemical pathways important in heat tolerance (48–50), and the effect of environmental factors such as use of sunscreen(51). We compare the findings of these studies to the observations in this work throughout our manuscript.

We hypothesized that metabolomes of coral species with different reported SCTLD susceptibility would vary in their metabolomes, and such variations could guide future work aimed at understanding of the pathways implicated in disease resilience. The sampling site in this study, Dry Tortugas, Florida, was being monitored regularly in anticipation of SCTLD beginning in September 2020, and the disease was first observed on May 29, 2021. Thus, the species in this study were not affected by SCTLD at the time of sampling (January 2020) but are representative of Florida species that are known to be susceptible to SCTLD. There are only a few investigations that compare metabolic or lipid profiles of field collected corals that are SCTLD susceptible(41). With the unabated spread of SCTLD along the Florida reef, opportunities to profile inter and intraspecies variation in metabolomes ahead of disease and post disease was envisioned to generate testable hypotheses to delineate biochemical pathways underlying disease susceptibility. To test our hypothesis and compare metabolomes of coral species with different SCTLD susceptibilities, we collected healthy coral fragments of four coral species, *Orbicella faveolata*, *Montastraea cavernosa*, *Meandrina meandrites*, and *Colpophyllia natans*, ahead of the SCTLD front in the Dry Tortugas. *M. meandrites* and *C. natans* are highly susceptible to SCTLD, while *O. faveolata* and *M. cavernosa* are defined as moderately susceptible (13, 15, 16). SCTLD susceptibility is defined by the length of time between the disease’s arrival to a reef and observation of lesions on a particular species, rates of lesion progression, and prevalence among species (15, 16). *M. meandrites* was one of the first reported species affected in the Dry Tortugas(52), while *C. natans* recruits spawned from parents in the Dry Tortugas showed *ex situ* lesion progression rates of 24.9-31.1%/day, which is in range for highly SCTLD susceptible corals (53). The Dry Tortugas is a unique habitat for corals along the Florida coral reef system where species such as *Acropora palmata*, *Siderastrea siderea,* and *Porites astreoides* have exhibited faster growth rates and enhanced reproduction relative to conspecifics in the Florida Keys (54). This may be attributed to oceanographic conditions that drive periodic upwelling, which is favorable for heterotrophy, and cooler temperatures and greater distance from urbanization and sources of pollution(54). Exogenous untargeted metabolome profiles of Dry Tortugas’ seawater samples were previously reported to be distinct from the profiles of seven other zones within Florida’s Coral Reef prior to SCTLD arriving at the Dry Tortugas(46). Thus, the visually healthy corals in this study from the Dry Tortugas provided the unique opportunity to examine and compare the metabolomes of several coral species growing under optimal growth conditions in Florida(54–56) before this region was affected by SCTLD.

In this work, we apply an untargeted high-performance liquid chromatography-mass spectrometry (LC-MS)-based approach to profile the metabolomes of a small sample set of four coral species (*M. meandrites*, *C. natans*, *O. faveolata*, *M. cavernosa*) utilizing recently developed advanced compound annotation methods to identify metabolites underlying the interspecies differences observed. While metabolomics analysis has been performed on Caribbean Scleractinian corals,(43, 45, 46, 57–61) our understanding of differences between the metabolomes of visually healthy Caribbean stony coral species is limited. Thus, we seek to address this gap in knowledge by describing chemical classes that are variably detected between four visually healthy coral species from the Dry Tortugas National Park sampled in January 2020 (52). In this study, we identify an endosymbiont-derived vitamin E pathway and a host-derived acylcarnitine pathway that were significantly variable among species. We describe additional chemical diversity by partitioning the crude extract of whole coral. Lastly, we report differences in the bioactivity of partitioned extracts of whole corals against bacterial pathogens.

## Results and Discussion

### Inter- and Intraspecies Variation in Metabolome Profiles

Metabolome extracts from four stony coral species (*Orbicella faveolata*, *Monstastraea cavernosa*, *Meandrina meandrites*, and *Colpophyllia natans*) were subjected to LC-MS analysis (Figure 1, Table S1). The resulting data was analyzed using a variety of data visualization and metabolite annotation tools (Figure 1C). Unsupervised principal component analysis (PCA) revealed that metabolome profiles of *M. meandrites* had the lowest intraspecies variation compared to the other coral species (Figure 2A). The largest interspecific separation captured on the first principal component (PC) was observed between *M. meandrites* and *M. cavernosa*. Four PCs captured interspecies variation between *M. meandrites* and the other species, and intraspecies variation for *M. cavernosa* (Figure S1A-E). The largest intraspecies distribution on PC1 was observed for *C. natans*, followed by *O. faveolata* and *M. cavernosa*. While the metabolomes of *M. cavernosa* fragments were spread across PC2, tighter clustering was observed for the other species along this component. PCs 3 and 4 captured metabolome variation between individual extracts, revealing that additional factors beyond species are captured within the metadata analysis (discussed further below) (Figure S1A-E). We calculated alpha and beta diversity for each coral (using Shannon entropy and Brays-Curtis similarity metrics, respectively, Figure S2) to quantify metabolome similarity. A principal coordinates analysis on the Bray-Cutis similarity matrix constructed on the metabolome data revealed tighter clustering for *M. meandrites,* while the other species were spread along the first principal coordinate (Figure S2A). The within species beta diversity was significantly larger than beta diversity between species (Figure S2B, Mann Whitney U Test, p=0.00082), further supporting that the metabolome analysis captures variation driven by factors beyond coral species. There was no significant difference between the alpha diversity for each species found using a Kruskal Wallis Test (Figure S2C, p= 0.119).

**Figure 1.**
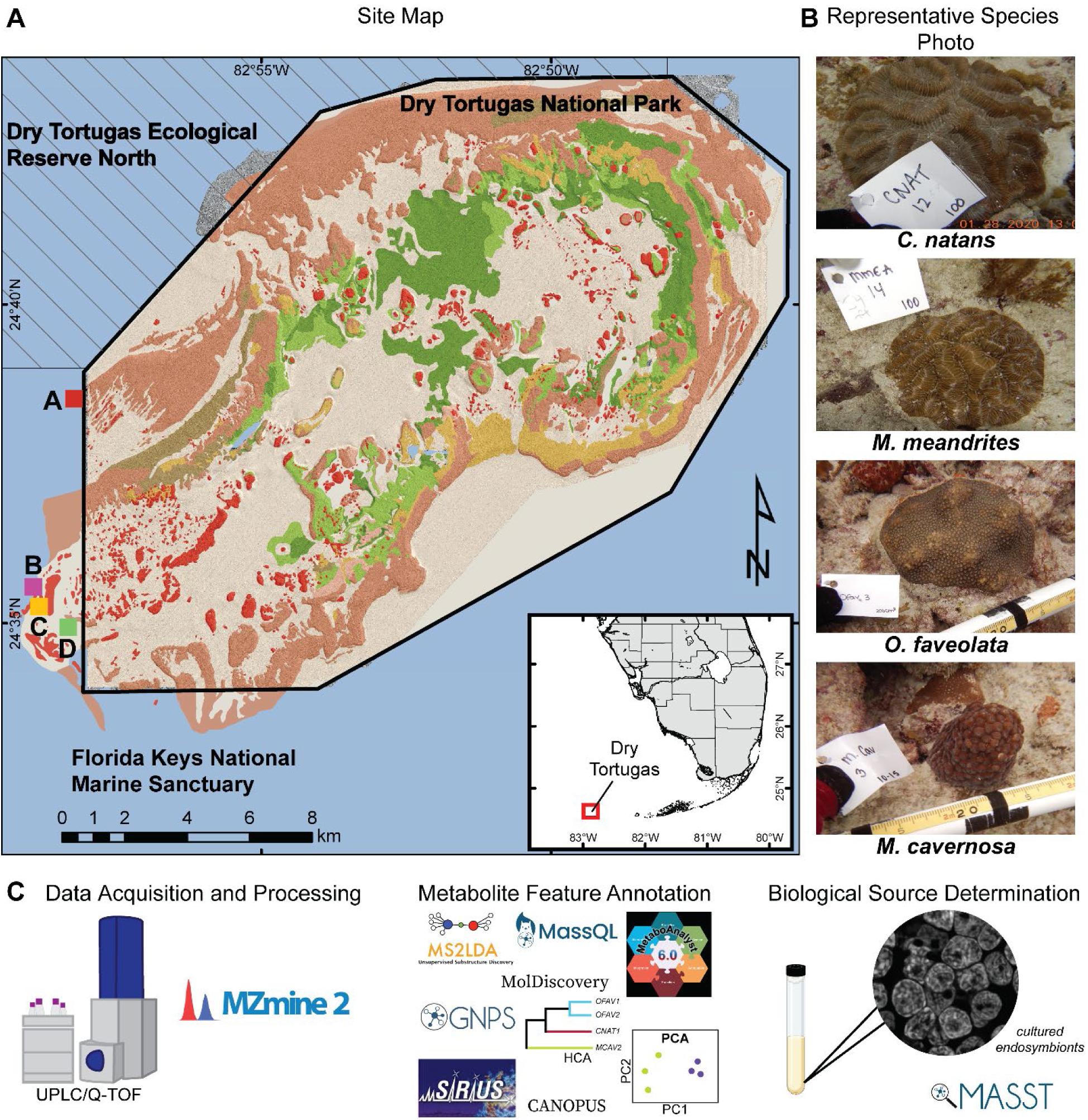
Sample collection, data acquisition, and analyses. **(A)** Benthic map of the sample sites in relation to Dry Tortugas National Park. Colors denote the benthic habitats (partially transparent) overlaid on high resolution bathymetry. Red and brown illustrates coral reef habitat (Florida unified reef map, Florida Fish and Wildlife Conservation Commission, 2016). A, B, C, D refer to the sites at which coral colonies were sampled. Coordinates for these sites are in Table S1. **(B)** Representative photographs of each coral species in this study (CNAT= *C. natans*, MMEA= *M. meandrites*, OFAV= *O. faveolata*, MCAV= *M. cavernosa*). **(C)** Untargeted metabolomics data were acquired, processed, and analyzed with a variety of methods. Metabolomics data available through public datasets (mined using MASST) and acquired on cultured algae was used to assign the biosynthetic source of annotated metabolite features.

**Figure 2.**
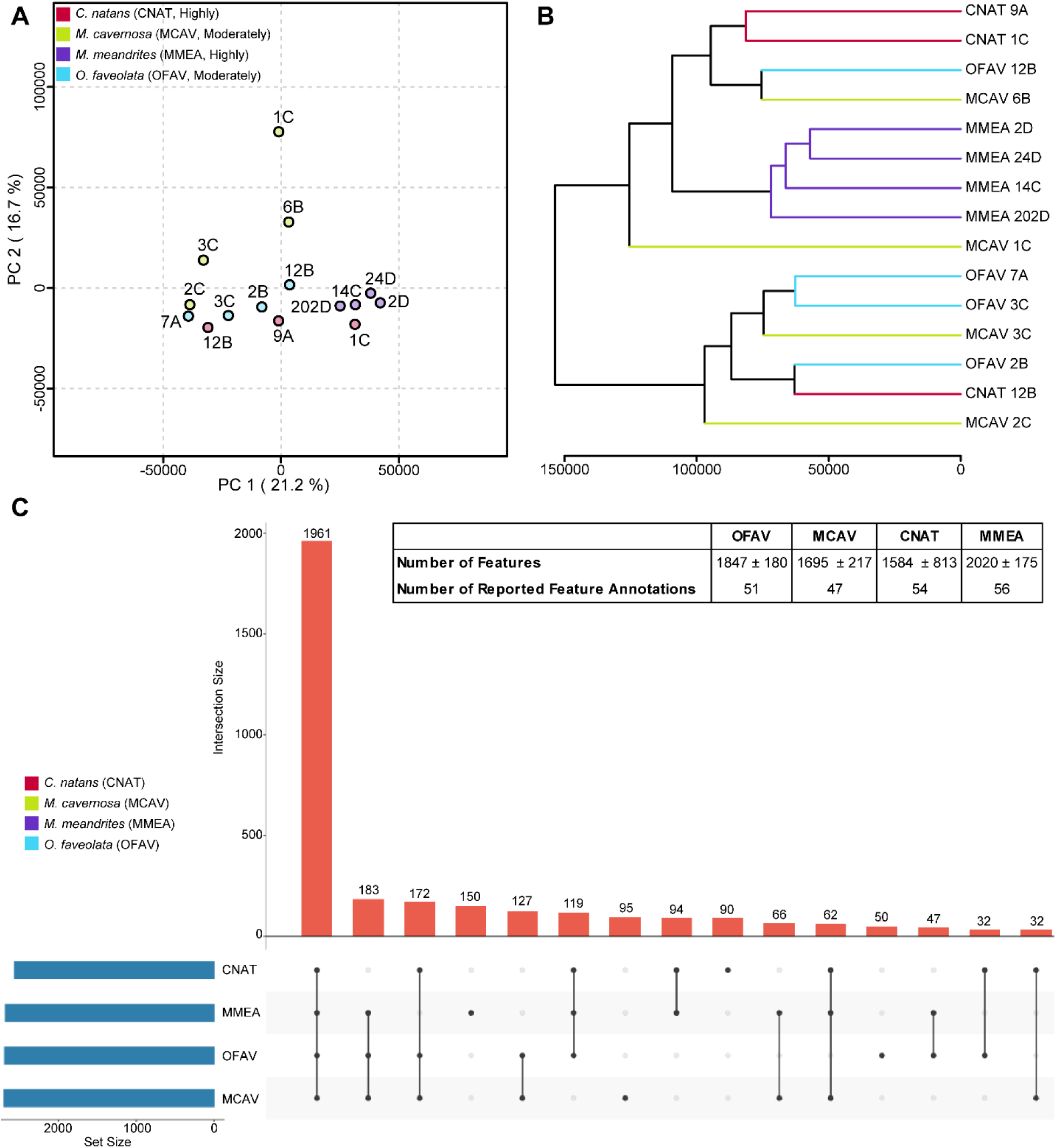
Intra- and interspecies variation of metabolomics data. **(A)** Principal component analysis of corals: *C. natans* (CNAT, red), *M. cavernosa* (MCAV, green), *M. meandrites* (MMEA, purple), and *O. faveolata* (OFAV, blue). The SCTLD susceptibility categorization is included in the key (‘Highly’, ‘Moderately’). Axes are labeled with the corresponding variance explained by each principal component. A, B, C, D refer to the site from which the coral was sampled (Table S1). **(B)** Hierarchical clustering analysis reveals a separate cluster for all *M. meandrites* samples, while the other species are distributed across clades. Colored branches correspond to species as outlined in (A). The letter at the end of the sample name corresponds to the sampling site. The x-axis represents the distance between the samples/clades. **(C)** UpSet Plot showing distribution of detected metabolite features. The number above each bar represents the number of features in that intersection. “Set Size” denotes the total number of features detected in each coral species. The inset table includes number of features detected within each species, as well as the number of annotated features reported in this manuscript.

Using permutational multivariate analysis of variance (PERMANOVA) (62), we found metabolome variation differed significantly across coral species (*Pseudo-F*_3,13_ = 2.192, p=0.007), with no significant effect of site (as a random effect, p=0.670). We queried whether SCTLD susceptibility categorization (OFAV, MCAV = moderate; CNAT, MMEA = high) affected metabolome variation. We could not examine interactive effects due to lack of Susceptibility x Species replication. We performed PERMANOVA with susceptibility as a single fixed factor and found that metabolome variation was significantly different between the two groups (*Pseudo-F* = 3.191, p=0.001). Additionally, a model with susceptibility as a fixed factor and species as a random effect found that Species(Susceptibility) was slightly significantly different (*Pseudo-F_1, 2_* = 1.715, p=0.021). Therefore, both coral species and SCTLD susceptibility affect metabolome variation, but it is unclear how the two interact in affecting metabolome differences. We cannot fully disentangle the effects of species and susceptibility with the current sampling regime in this study, therefore such efforts are an important avenue for future inquiry. Sampling species with moderate resilience (i.e. some individuals are susceptible, others never develop lesions) where both affected and unaffected colonies were sampled would be a potential way to disentangle this effect. A non-metric multi-dimensional scaling (nMDS) plot by species was constructed to visualize metabolome variation using a method appropriate for smaller sample sizes. The nMDS plot divided the samples into four distinct clusters. Consistent with the PCA, all *M. meandrites* samples clustered together in one group, and a larger spread was observed for other species with metabolomes of *M. cavernosa* displaying the largest intraspecies variation (Figure S3). These observations aligned with previous findings where apparently healthy *M. cavernosa* from a SCTLD endemic site in Broward County showed similar intraspecies variation(43). The proximity between *M. cavernosa* colonies on the reef in this previous study explained the variation in some instances but did not completely explain the metabolome variation observed for *M. cavernosa* in Broward County(43). The beta diversity analysis conducted in this study revealed three pairs of corals from the site B or D as having the greatest Bray-Curtis similarity score to each other. These include OFAV2B/CNAT12B, MCAV6B/OFAV12B, and MMEA2D/MMEA24D. Thus, we see further evidence of reef site driving metabolome similarity in some instances, although for the MMEA pair we cannot disentangle the effect of site and species on the metabolome similarity.

An unsupervised hierarchical clustering analysis (HCA) revealed *M. meandrites* was the only species that clustered within a single clade (Figure 2B, purple branches), further indicating the relatively low intraspecies variation. The HCA also reveals that the corals within the aforementioned pairs identified through the Bray-Curtis similarity analysis have the greatest metabolome similarity to each other (Figure 2B). There was significant variation in metabolomic variation (multivariate dispersion) among the coral species (F_1,3_=7.944, p=0.045). Overall, MCAV was the most variable (average distance to group centroid = 33.3) with the variation being significantly higher than the variation observed for MMEA (average distance = 19.2, p = 0.036). OFAV was the second most variable (average distance = 29.3) and the variation was significantly higher than the variation of MMEA (p=0.030). The relative metabolomic variation did not differ between pairwise comparisons performed for other coral species. This phenomenon has been observed in deep sea corals where interspecies differences rather than site dictated clustering of metabolite profiles(63). However, intraspecies differences may be attributable partially to site, as Haydon and colleagues found metabolite differences in *Pocillopora acuta* based on reef site, even after acclimation of the corals in aquaria as is the case in this study(48). Thus, when comparing multiple species of corals from different reef sites, replication of species collected from each site should be conducted where possible. The intraspecies metabolomic variation observed in this study may further be partially explained by different genotypes(45), and both intra- and interspecies variation may be influenced by the endosymbiotic profile, microbial community, bleaching history, and stimuli/stressors unique to the sampling site(21, 22, 64, 65). Acquiring data on seawater (exometabolomics), cataloguing abiotic factors at reef sites, and profiling the endosymbiont and microbial community will aid in disentangling what additional factors drive metabolome variation. In a metabolomics study of cultured *Symbiodinium* species, Klueter *et al* noted that metabolite profiles varied by species and the degree of metabolome variation was not ubiquitous across species given the different classification error rates of each symbiont species(66). The distinct metabolome profiles of *M. meandrites* compared to the other corals species in this study could be influenced by the symbiont types of the coral species. The *M. meandrites* sampled for this study may host symbionts with highly similar metabolomes or interaction networks with the associated microbiome, while the other coral species may host symbionts and/or microbiome with more diverse metabolomes. Incorporating microbiome analysis and symbiont typing into metabolome studies would be beneficial towards delineating how the degree of observed metabolome variation may correspond to the symbiont and the microbiome species present. Another factor that may contribute to the intraspecies metabolome variation captured in this study is cryptic lineages observed within the studied species(67). Cryptic coral species lineages may share phenotypic traits but have distinct underlying genomic differences(67). *M. cavernosa*(68, 69) and *O. faveolata*(70, 71) are reported to have cryptic lineages. The genomic differences between the coral hosts imply a strong possibility for metabolite differences (since the metabolome reflects the functional biochemical state of the system, in this case the coral holobiont). Proven association with different symbiont genera (reported for *O. faveolata*(70)) and potential association with different microbial assemblages (yet to be studied for the species in this report) among cryptic lineages could further diverge metabolome profiles, as all the members of the coral holobiont influence and contribute to the metabolome. Such phenomena(67) may well explain the metabolome variation of *M. cavernosa* and *O. faveolata* in this study, and although cryptic lineages have not yet been identified for *Colpophyllia* (67) the observed metabolome variation in *C. natans* may be partially explained by this as well.

The UpSet Plot generated to show the distribution of metabolite features revealed the greatest number of unique features were detected in *M. meandrites* extracts, followed by *M. cavernosa*, *C. natans*, and *O. faveolata* (Figure 2C). Since the statistical analyses revealed a distinct metabolome profile for *M. meandrites* and a greater intraspecies metabolite variation captured for the other coral species, unique features present in *M. meandrites* and features that were variably detected among *M. meandrites* and other coral species were prioritized for annotation and are described below.

### Metabolite Features Driving Variation

#### Vitamin E family compounds as potential biomarkers of stressor susceptibility

A metabolite feature *m/z*_RT (*m/z*: mass to charge, RT: retention time in min) 449.398_21.3 min, uniquely detected in *M. meandrites* extracts, was proposed as α-tocopherolhydroquinone by the in silico annotation tool MolDiscovery. SIRIUS with CSI:FingerID also proposed the annotation for this feature as α-tocopherolhydroquinone. We searched for α-tocopherolhydroquinone spectra in the literature and used MS^2^ spectral matching with a published spectrum(72) of silicated α-tocopherolhydroquinone to further support the annotation (Figure S4A). This feature clustered in Feature Based Molecular Networking (FBMN) analysis with another feature at 447.383_21.3 min representing one unsaturation from α-tocopherolhydroquinone (Δ*m/z=* 2.015). We annotated this feature as α-tocopherolquinone, the oxidation product of α-tocopherolhydroquinone(73, 74) and confirmed this annotation with an analytical standard of α-tocopherolquinone (Figure S4B-C). To identify additional metabolites we applied unsupervised substructure discovery using MS2LDA(75) (Figure 3A, Table S2).

**Figure 3.**
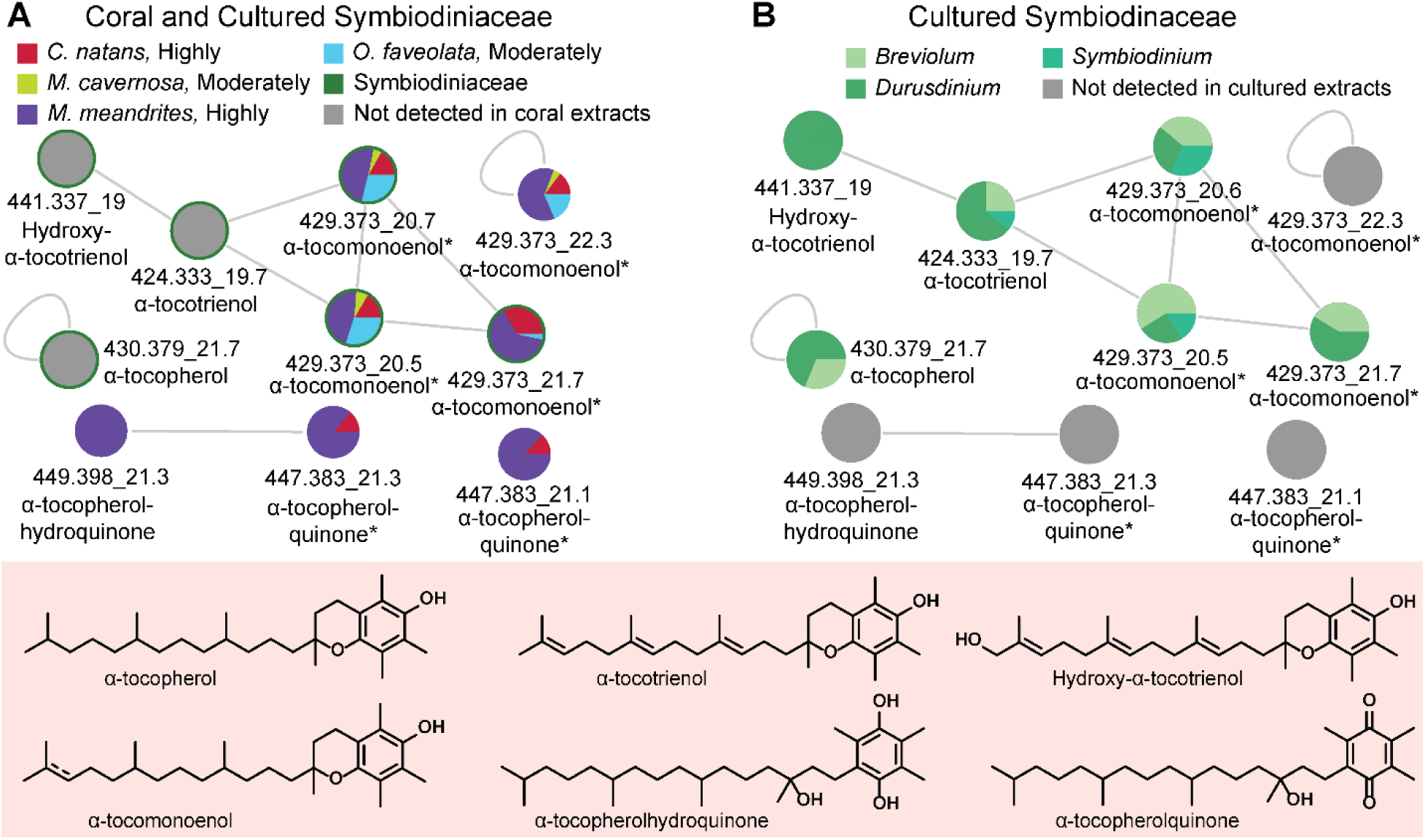
Analyses of vitamin E family compounds. **(A)** Network of annotated vitamin E family compounds with *m/z*_RT (*m/z*: mass to charge, RT: retention time in min). The * indicates these features have identical MS^2^ spectra, but different retention times, representing isomeric species. **(B)** The detection pattern of annotated vitamin E family compounds in cultured endosymbiont extracts. The comprehensive list for annotated vitamin E family compounds is provided in Table S2, and MS^2^ mirror plots supporting annotations of tocopherol(hydro)quinone are provided in Figure S4. The SCTLD susceptibility categorization is included in the key (‘Highly’, ‘Moderately’).

It is interesting to note that α-tocopherolquinone, an oxidation product of α-tocopherol(76, 77), and α-tocopherolhydroquinone were exclusively detected in highly SCTLD susceptible *M. meandrites* and *C. natans*, but not in *O. faveolata* and *M. cavernosa* with moderate SCTLD susceptibility(15, 16). These features were also not detected in the extracts of cultured Symbiodiniaceae included in this study (Figure 3B). The *Durusdinium-*associated mangrove coral *Pocillopora acuta* was previously reported by Haydon and colleagues to accumulate α-tocopherol in summer(48). This observation was suggested as a possible mechanism of ‘frontloading’ associated with the resilience of *Durusdinium*-associated corals (48). Tocopherols are the most prominent antioxidants that counteract lipid peroxidation. Using transcriptomics, the gene for arachidonate 5-lipoxygenase (ALOX5), was found to be significantly differentially expressed with its highest expression in the most susceptible corals, including *C natans*(39). ALOX5 is a key enzyme in mediating lipid peroxidation(78) which can lead to cell death such as apoptosis, ferroptosis, and pyroptosis(78). Thus, we searched the literature to identify studies that might link α-tocopherol(hydro)quinones with lipid peroxidation and cell death. While linking specific metabolites with processes is outside the scope of this study, we can speculate on possible functions of metabolites and determine future avenues of inquiry based on literature precedence. Indeed, a recent report updated the mechanism-of-action for iron-dependent anti-apoptotic activity (ferroptosis) of α-tocopherol(79). Tocopherol was suggested to be the pro-vitamin E form, while the (hydro)quinone forms produced from oxidation of α-tocopherol were shown to be the activated forms responsible for prevention of cell death(79). Thus, it is possible that our detection of the α-tocopherol(hydro)quinones indicates the coral cells are frontloading the activated form of vitamin E, which is counteracting lipid peroxidation resulting in detection of α-tocopherol(hydro)quinones.

When corals were previously challenged with bacterial pathogen-associated lipopolysaccharides, susceptible corals demonstrated a transcriptome response related to apoptosis, while resistant corals transcribed genes related to autophagy, a more modulated response to stressors(80). The damage threshold hypothesis proposes that coral disease susceptibility is inversely related to the upper limit of damage a coral can withstand before harmful effects are observed(81). Corals with a low damage threshold (high susceptibility) may be unable to modulate immune responses; either mounting too high of a response, leading to auto-immune challenges, or too low of a response before cellular death is imminent. Thus, the varied detection of tocopherol(hydro)quinones in this study should be further investigated to determine if these metabolites serve as a biomarker of corals particularly susceptible to disease, represent stressor history, and if they vary temporally with disease progression.

*M. meandrites* and *O. faveolata* had the highest relative abundance of α-tocomonoenol (Figure 3A). α-tocomonoenol was previously detected at higher abundance in apparently healthy *M. cavernosa* compared with diseased corals(43). The analogue α-tocotrienol (*m/z* 424.333) with three degrees of unsaturation, was exclusively detected in SCTLD-affected *M. cavernosa*, while other unsaturated analogs were likewise detected at higher abundance in the diseased corals(43). In this study, where we have analyzed healthy corals ahead of the SCTLD front, α-tocotrienol was not detected. Based on these results, we hypothesize that tocotrienols may serve as biomarkers for coral disease, wherein accumulation coincides with disease progression. A time course study that tracks how tocotrienol analogues and tocopherolquinones accumulate in response to disease exposure is required to validate this hypothesis. Recent work highlighting differential detection of tocopherol upon heat stress among resilient and susceptible species(48, 82) and our work reporting detection of different tocopherol analogues among healthy and SCTLD-affected coral colonies suggest that this endosymbiont pathway likely plays an important role in coral health and resilience(48, 83); warranting studies that monitor tocopherol-related metabolite production over time after disease exposure.

### Acylcarnitine profiles differentiate *Meandrina meandrites*

Feature 476.373_14.4 min was proposed by SIRIUS with CSI:FingerID as docosatetraenoyl carnitine (C22:4) (Figure 4A, B). To confirm this annotation prediction and to determine if other acylcarnitines were present in our data, the output of the MS2LDA analysis was consulted. This feature shares MS2LDA substructure motif 185 with feature 276.180_2.8 min, which was annotated as hydroxyhexanoyl carnitine (C6:0-OH) based on the MS^2^ fragment peak at *m/z* 217.107 associated with the neutral loss of trimethylamine (*Δm/z=* 59.07, Figure 4C, S4A). A variety of acylcarnitines were further annotated with the aid of the GNPS spectral library, MassQL, substructure motif 185, and SIRIUS with CSI:FingerID (Figure 4, S4B-O, Table S2).

**Figure 4.**
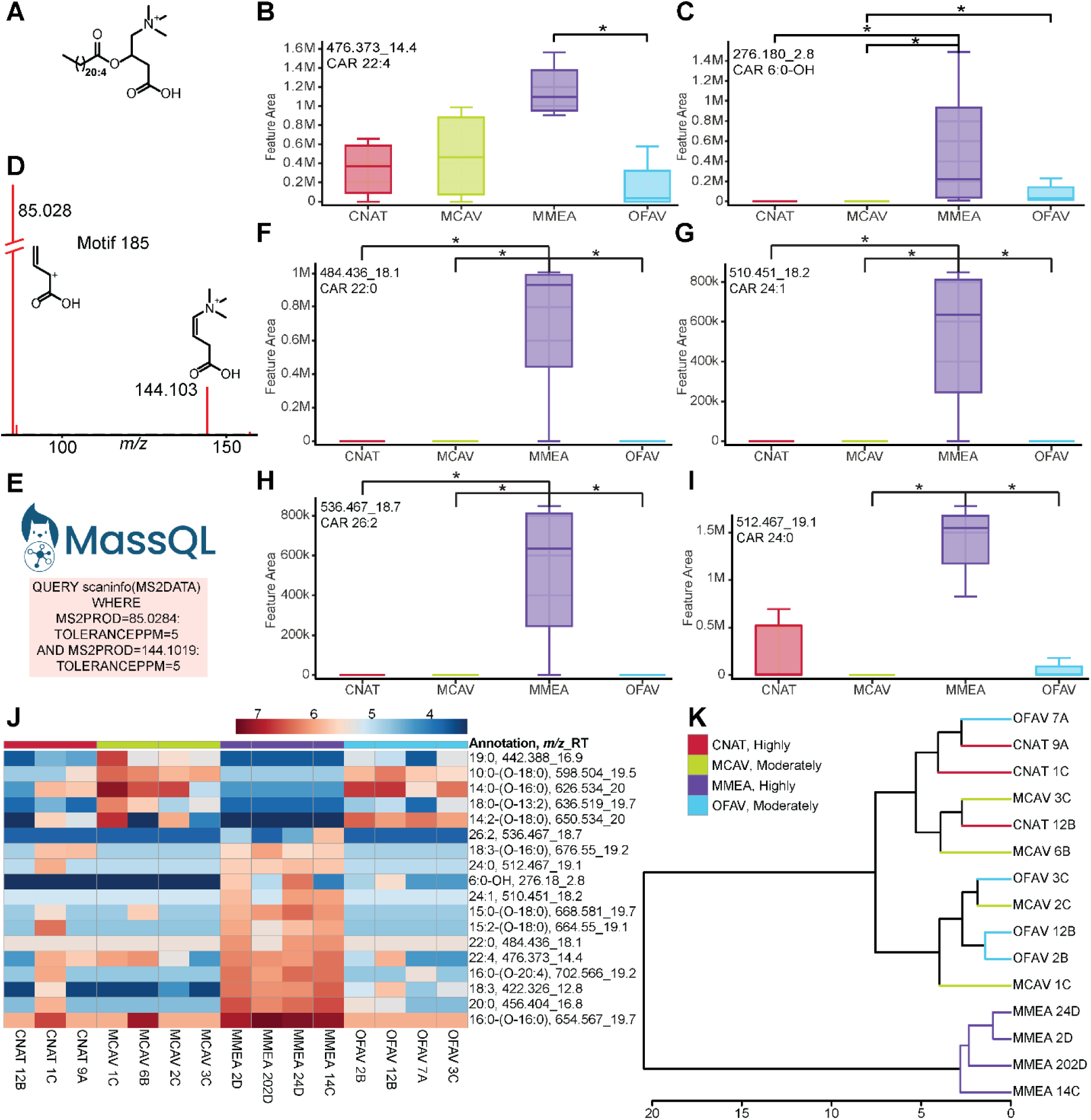
Annotation of and variation in detected acylcarnitines. **(A)** Chemical structure of docosatetraenoyl carnitine (C22:4). **(B, C, F-I)** A subset of the box plots of acylcarnitines differentially detected in the extracts of *M. meandrites*. Asterisks indicate significant differences as determined by a Kruskal Wallis test with Dunn’s post-hoc test (adjusted *p*<0.05). Additional box plots are reported in Figure S4. **(D)** MS2LDA motif 185 used to aid annotations of features annotated as acylcarnitines. **(E)** The query submitted to MassQL to search for acylcarnitines. **(J)** Heat map of the features annotated as acylcarnitines that were determined as statistically differentiating by Kruskal Wallis test with Dunn’s post-hoc test (adjusted *p*<0.05). The scale bar represents the log-transformed abundance. The putative annotation and *m/z*_RT are included for each feature (Table S2 and Figure S5). **(K)** The hierarchical clustering analysis based on the log-transformed abundances of the acylcarnitines. The SCTLD susceptibility categorization is included in the key (‘Highly’, ‘Moderately’).

Acylcarnitines are typically host-derived metabolites, and these metabolites were not detected in the cultured Symbiodiniaceae extracts in this study. Acylcarnitines have been detected at higher abundances in the daytime exometabolomes of *Porites* and *Pocillopora* compared to algae (turfing microalgae, macroalgae, and crustose coralline algae), where they are hypothesized to play a role in nitrogen and phosphorous cycling(84). Acylcarnitines play an integral role in metabolism of fatty acids in mitochondria(85) and maintenance of available pools of free coenzyme A(86). In the diatom *Phaeodactylum tricornutum*, propanoyl-carnitine and butanoyl-carnitine accumulate under nitrogen-starvation(87). Accumulation of acylcarnitine concentrations have been linked with cell toxicity(88), mitochondrial dysfunction(89–91), and dysfunction in cellular bioenergetics in humans(88). Acylcarnitines have been found to be upregulated in corals upon exposure to octocrylene, an ingredient used in sunscreens(51), which is the only study reporting conditional dysregulation of acylcarnitine levels in corals found in our literature search. In this study, the interspecies variation of all features annotated as acylcarnitines were analyzed with a Kruskal Wallis test with Dunn’s post-test (adjusted *p*<0.05). The acylcarnitines with fatty acyl tails with C13-C20 are classified as long chain, and tails >C21 as very long chain(92). Features that were differentially detected showed two interspecies patterns based on the acyl chain length (Figure 4B, C, F-J and S4D-O). The hydroxyhexanoyl acylcarnitine and very long chain acylcarnitines were detected at higher intensity in *M. meandrites* (Figure 4 B, C, F-I). The accumulation of long chain acylcarnitines is associated with several metabolic diseases in humans(92). Differences in acylcarnitine profiles have also been reported as indicators of frailty in humans(93). Given that certain acylcarnitine analogues are detected at higher intensity in *M. meandrites* (Figure 4 and S4), a highly SCTLD-susceptible species, it is possible that acylcarnitine profiling could represent disease history and/or higher susceptibility to disease. Interestingly, when an HCA was performed on only the annotated acylcarnitine features, a clear separation of *M. meandrites* from other coral species was observed (Figure 4K). Thus, host-derived acylcarnitines display a species-specific profile. Since several acylcarnitines were variably detected in these apparently healthy corals, the understudied role of carnitines in disease resilience and susceptibility in corals should be further investigated.

Several unknown acylcarnitines were distributed differentially across the coral species (Figure 4J and S4G-O). Upon manual inspection of fragmentation spectra, we propose the annotation of these features as acylcarnitines containing fatty acid esters of hydroxy fatty acids (known as FAHFAs) (Figure S5A, Table S2). The fragments at *m/z* 85.028 and 144.102, presence of a fragment corresponding to hydroxylated fatty acid of acylcarnitine (CAR 14:1-OH), and the presence of an additional fatty acid tail fragment (C18:0) supported the annotation of an FAHFA-containing acylcarnitines (Figure S5A, bottom spectrum; CAR 14:1-(O-18:0)). FAHFAs are a conserved class of lipids that are widely reported, including in dietary plants(94, 95), as defense molecules in caterpillars named as mayolenes (96, 97), as anti-inflammatory metabolites in humans(98), and in the corallivore Crown-of-Thorns Starfish(99). Oxidative and environmental stress increase the synthesis of FAHFAs and ornithine-conjugated FAHFAs(100, 101). Acylcarnitines containing FAHFAs have not been previously reported and warrant further investigation for structural characterization and their role in coral biology. We searched for these acylcarnitines features in the publicly available datasets on the MassIVE server using MASST(102). These features were found in several marine organism-derived datasets including datasets from several coral species (Table S3) but were not observed in human-derived datasets. These observations further strengthen the role and application of modern methods in data analysis in untargeted metabolomics to discover biologically relevant metabolic pathways and generate testable hypotheses. Here, access to public datasets on these pristine endangered coral species is advantageous.

### DGCC betaine lipids with 16:0 fatty acyl tails are differentially detected between species

Several differentiating features were identified as diacylglyceryl-carboxyhydroxymethylcholine (DGCC) betaine lipids. Feature 774.584_19.6 min was a GNPS library match to DGCC(36:5) (Figure 5A, Figure S6A). The fragment peaks at *m/z* 490.373 and 472.363 in the MS^2^ spectra, which are characteristic of the chemical substructure containing a 16:0 fatty acyl tail, enable further annotation of this feature as DGCC(16:0_20:5) (Figure S7). This feature was variably detected among coral species, present at highest abundance in *M. cavernosa*. As expected, the feature 490.373_13.1 min, annotated as lyso-DGCC(16:0) known to be a constituent of healthy corals(43, 58, 103), was detected in all species (Figure 5B, Figure S6B). We used MassQL to search for additional DGCC analogues containing a 16:0 fatty acyl tail (Figure S6C). This approach permitted the annotation of additional metabolite features, detected at highest abundances in *M. meandrites*, as lyso-DGCC(16:0) analogues (Figure 5 and S7, Table S2). Diacylated and unsaturated DGCC betaine lipids have been previously proposed as biomarkers of coral bleaching(58, 103). The increase in lipid unsaturation is suggestive of increased cell death when the antioxidative capacity of cells is lowered(104). DGCC betaine lipids are biosynthesized by Symbiodiniaceae(50). We searched the metabolite data acquired on cultured Symbiodiniaceae for the presence of the annotated DGCC analogues. While monoacylated lyso-DGCC(16:0) was detected in all cultured Symbiodiniaceae genera, the diacylated analogues were notably absent in *Durusdinium* extracts, the genera known to be most thermotolerant(105–108) (Figure 5H). Roach *et al*. noted a higher abundance of lyso-DGCCs in historically non-bleached corals, while unsaturated and DGCCs were abundant in historically bleached corals(58). Rosset *et al*. observed significantly higher abundance of lyso-DGCC and unsaturated DGCCs in thermotolerant *D. trenchii* as compared to *Cladocopium* C3 in both control and heat stressed conditions(49, 50). Symbiodiniaceae genera show differential responses to thermal and irradiance stress, which affects the entire holobiont response to stressors(64, 109–111). The variable detection of the DGCC(16:0) analogues in the coral extracts may indicate variable bleaching history or the presence or absence of certain Symbiodiniaceae species in the coral colonies sampled. Previous reports suggest that the DGCC lipid profile is influenced by the host(112). Given that algae transform their membranes in response to a variety of stimuli and stressors(113–115), it is also possible that the variable detection of the diacylated DGCC(16:0) analogues is reflective of host-dependent shifts in betaine lipid profiles of Symbiodiniaceae.

**Figure 5.**
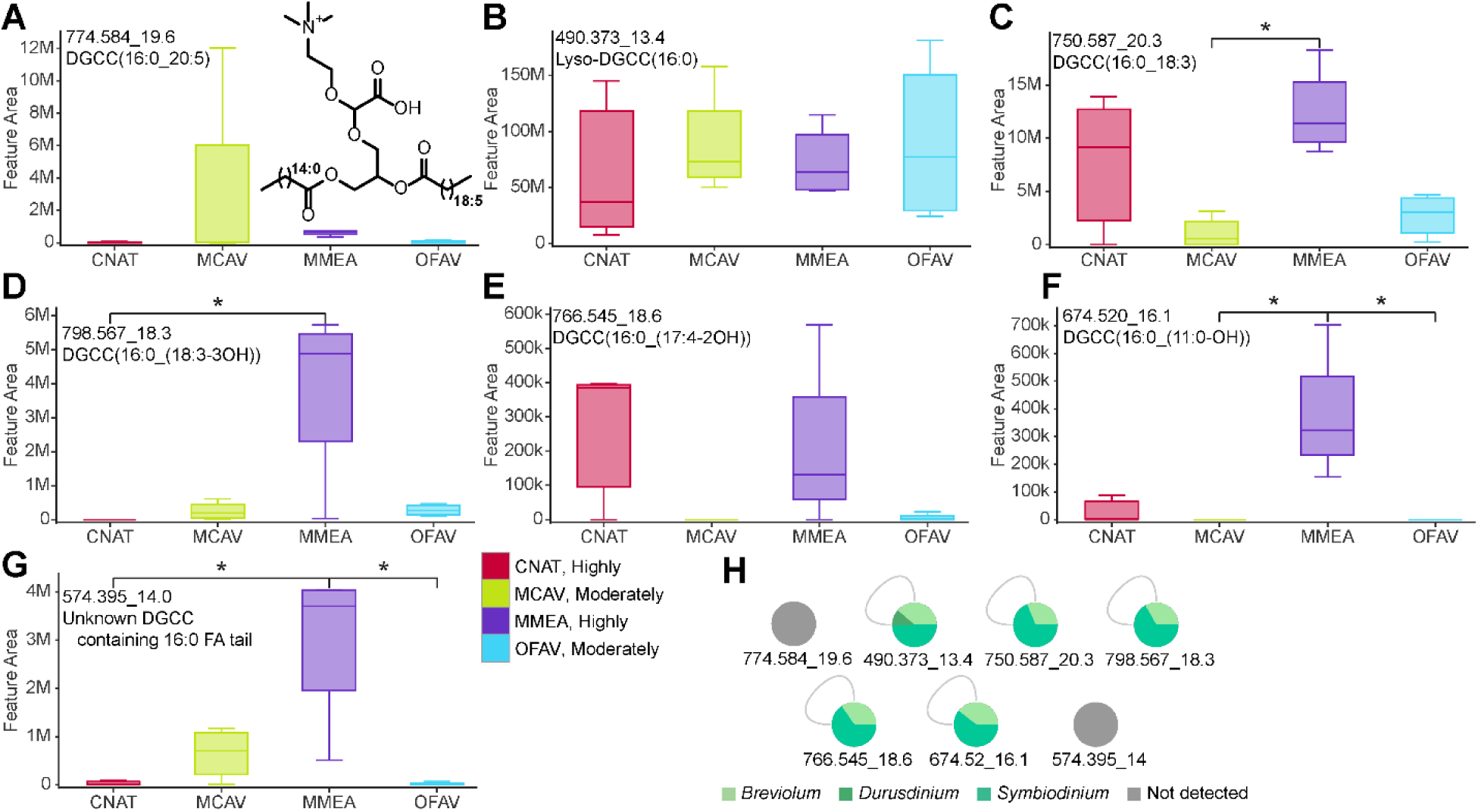
Interspecies variation of diacylglyceryl-carboxyhydroxymethylcholine (DGCC) betaine lipids. **(A-G)** Box plots of features annotated as DGCC(16:0) analogues. Annotations are provided in Figure S6, S7 and Table S2. Asterisks indicate significant differences between coral species as determined by Kruskal Wallis test with Dunn’s post-hoc test (adjusted *p*<0.05). **(H)** The detection pattern of DGCC analogues in the cultured zooxanthellae extracts. The SCTLD susceptibility categorization is included in the key (‘Highly’, ‘Moderately’).

### Carotenoid pigments do not show coral species-specific patterns

Carotenoids are important antioxidants in photosynthetic organisms. Symbiodiniaceae produce several carotenoids such as peridinin, fucoxanthin, astaxanthin, diatoxanthin, diadinoxanthin, and neoxanthin, with peridinin being the most prevalent and abundant(116, 117). We examined whether endosymbiont-derived pigment profiles contributed to variation among the coral colonies analyzed in this study. Several pigments were annotated using mass spectral search and literature search (Figure 6 and S8A-F, Table S2). The features annotated as pigments were also analyzed by HCA (Figure 6B). The pigment profile did not display clear interspecies variation but did display intraspecies variation. Peridinin was detected in all coral extracts, while fucoxanthin was detected in only a few coral extracts (Figure 6A). Among cultured Symbiodiniaceae in this study, peridinin was detected in all genera whereas fucoxanthin was only detected in thermotolerant *Durusdinium* cultures (Figure S8G). Since Wakahama *et al*. reported a negative correlation between presence of fucoxanthin and peridinin in a variety of symbiotic and free living *Symbiodinium* strains(118), we confirmed detection of fucoxanthin in peridinin-containing coral extracts using an analytical standard of fucoxanthin (Figure S8F). Detection of both pigments may represent presence of multiple strains of Symbiodiniaceae within the cultures.

**Figure 6.**
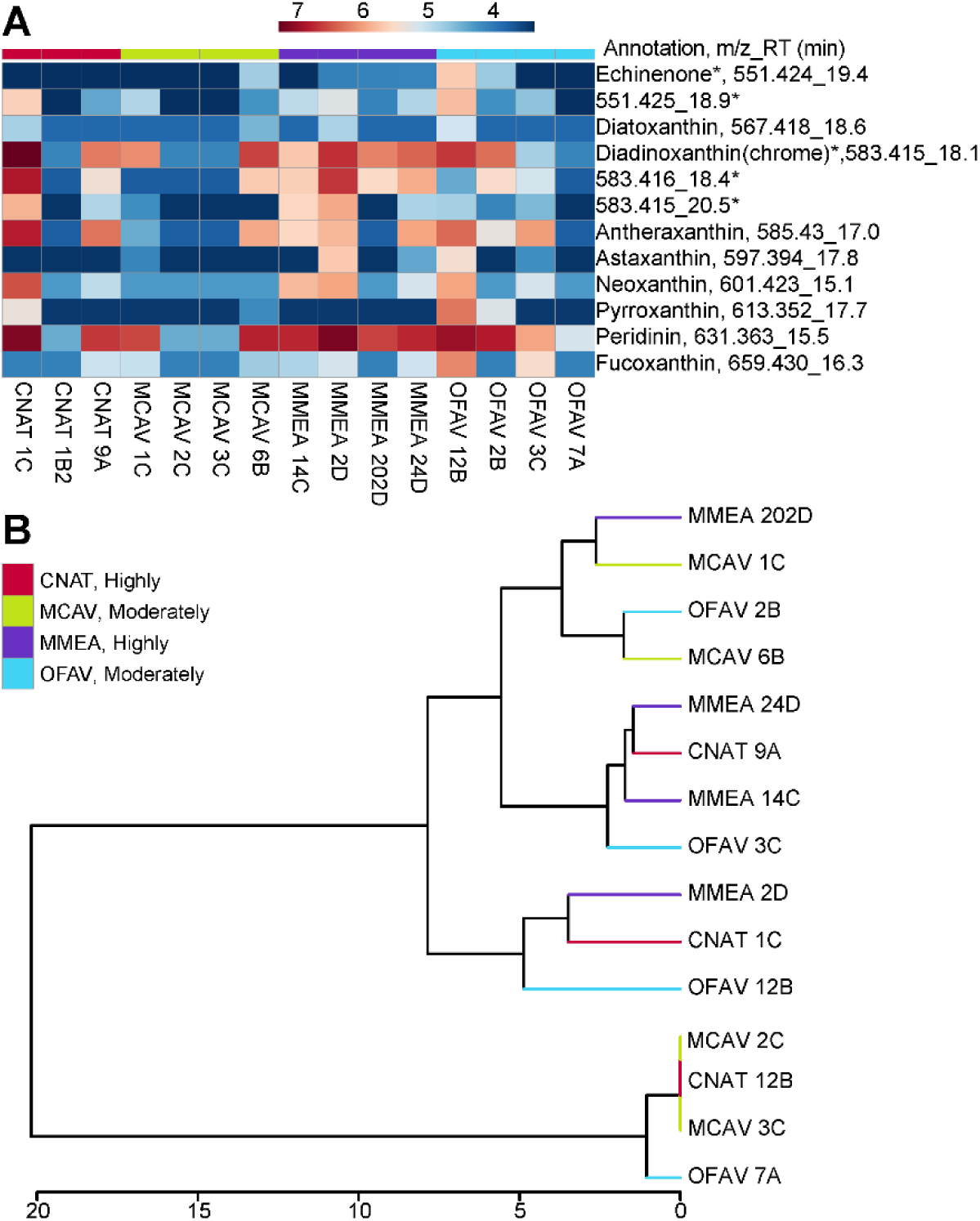
Analyses of endosymbiont-derived pigments. **(A)** Heat map showing the distribution of features annotated as pigments. The log-transformed abundance is reported. The *m/z*_RT and annotation are included. The * indicates these features have identical MS^2^ spectra, but different retention times representing isomeric species **(B)** Hierarchical clustering analysis based on the log-transformed abundance of the annotated pigments shows no clustering by species nor site. The SCTLD susceptibility categorization is included in the key (‘Highly’, ‘Moderately’).

### Butanol partitions of whole coral extracts enable additional metabolite annotations

The crude extracts from the whole coral samples were further partitioned into ethyl acetate (EtOAc) and butanol (BuOH) solvents and the bioactivity of these fractions was tested against the potential SCTLD-associated pathogens *Vibrio coralliilyticus* OfT6-21 and OfT7-21, *Leisingera* sp. McT4−56, and *Alteromonas* sp. McT4-15(119, 120) using an agar disk-diffusion assay (Figure 7A, Figure S9A). Partitions only exhibited activity against *V. coralliilyticus* strains. The largest zones of inhibition were observed for BuOH partitions of *M. meandrites* against both pathogens (Figure 7A and S9A). Thus, untargeted metabolomics data were acquired on BuOH partitions of all species. The metabolite data was analyzed following the scheme outlined in Figure 1. Within the BuOH partitions, an additional 560 metabolite features were detected (Figure 7B). The UpSet Plot analysis showed the greatest number of unique features was detected in *C. natans* extracts, followed by *M. cavernosa*, *M. meandrites*, and *O. faveolata* (Figure 7B). We used CANOPUS to predict the chemical classes of these features (Table S4). For the features uniquely detected in the BuOH partitions, none of the metabolites in the CANOPUS-predicted natural product pathways were significantly enriched in *M. meandrites* compared to the other species (Figure 7C). The UpSet Plot and CANOPUS output were used to guide compound annotations (Figure 7D).

**Figure 7.**
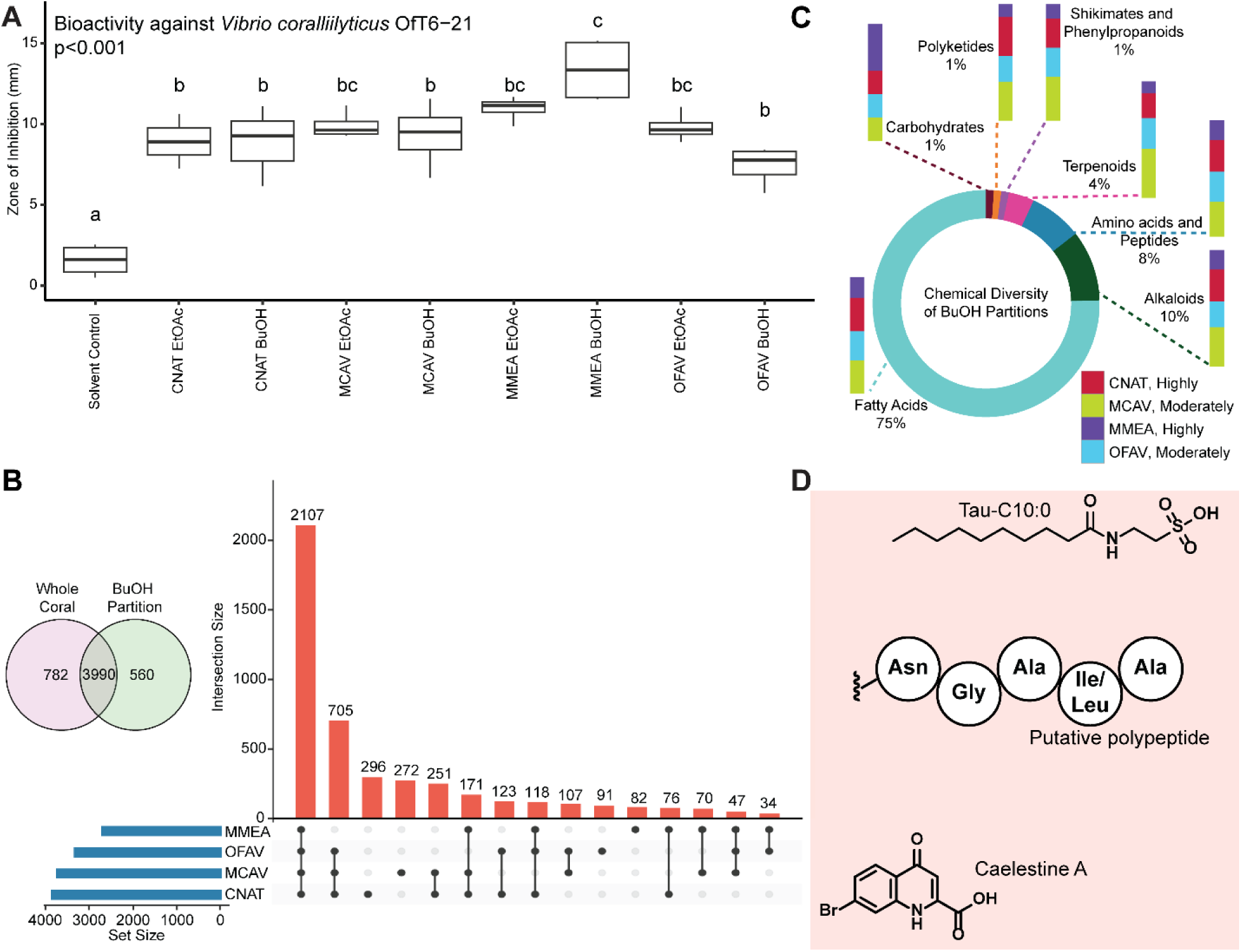
Analysis of BuOH partitions of crude extracts. **(A)** Bioactivity of EtOAc and BuOH partitions of crude extracts determined using agar diffusion growth inhibition assay against coral pathogen *V. coralliilyticus* OfT6-21. Letters denote results from a Tukey HSD posthoc test following a significant one-way ANOVA (p<0.001). **(B)** Venn Diagram of features detected in whole coral extracts and BuOH partitions. 560 unique metabolite features were detected in BuOH partitions. UpSet Plot of all features detected in BuOH partitions, where the number above each bar represents the number of features in that intersection and “Set Size” denotes the total number of features detected within that coral species. (MMEA= *M. meandrites*, OFAV= *O. faveolata*, MCAV= *M. cavernosa*, CNAT= *C. natans*). The SCTLD susceptibility categorization is included in the key (‘Highly’, ‘Moderately’). **(C)** Chemical diversity of features detected exclusively in BuOH partitions, along with the distribution of these features among the coral species. Natural product pathways (probability >0.8) were determined using CANOPUS. **(D)** Chemical structures of putatively annotated metabolite features uniquely detected in BuOH partitions (annotations supported in Figures S8 and S9).

A feature, detected exclusively in BuOH partitions at 280.157_7.9 min was annotated as Tau-C10:0 based on MS^2^ spectral matching (Figure S10A). We also observed the presence of the *N*-acyl taurines in several publicly available datasets acquired on diatoms, dinoflagellates, and seawater by searching the MS^2^ spectrum of this metabolite in MASST (Table S3). *N*-acyl taurines have been implicated as important signaling molecules in several human processes including postprandial glucose regulation(121), but these molecules have not been previously described in corals. Thus, partitioning crude extracts into organic solvents can enable detection and characterization of low-abundance metabolites, which are otherwise below the limit of detection. Feature 267.960_5.3 min with an isotopic pattern of a brominated compound was uniquely detected in the BuOH partitions of *O. faveolata*. This annotated as caelestine A based on MS^2^ spectral matching and MS^1^ isotopic pattern (Figure S10B-C). CANOPUS predicted the chemical class as hydropyrimidine carboxylic acids and derivatives. Caelestine A, a brominated quinoline carboxylic acid, has been reported as a possible indicator of a response to heat stress in the invasive bryozoan *Bugula neritina*(122). A feature at 615.346_5.9 min was proposed as a tunicyclin G analogue by DEREPLICATOR, which compares experimental MS^2^ spectra against predicted in silico MS^2^ spectra of peptides(123) and CANOPUS predicted the chemical class of this feature as oligopeptides. We putatively annotated the feature 615.346_5.9 min as a polypeptide with partial sequence NGAI/LA (Figure S9C). This polypeptide was detected in *M. cavernosa* and *O. faveolata* BuOH partitions. Although we could not link the enhanced antibacterial activity with specific molecules or chemical classes due to a small sample size, these annotations show that partitioning of whole coral metabolomic extracts increases the breadth of detected metabolites and enables annotation of low-abundance natural products. Annotation of individual metabolites is still a tedious and manual task. As spectral libraries are populated by the community, and in silico compound annotation methods advance, these datasets will be a valuable resource to untangle mechanisms of symbiosis between members of coral holobionts.

### Conclusion

The visually healthy corals collected from the Dry Tortugas, a region of Florida with oceanographic conditions that support coral productivity, revealed interspecies metabolome differences. Tight clustering of the *M. meandrites* metabolome indicated similar metabolite profiles, while higher metabolome variation was found for *C. natans*, *M. cavernosa*, and *O. faveolata*. Metabolites driving the variation between species included tocopherol(hydro)quinones, diacylated betaine lipids, and acylcarnitines. This is the first report describing differences in acylcarnitine profiles between coral species and the discovery of potentially novel analogues containing an additional fatty acid group. Given the specificity of these acylcarnitine-FAHFA to only marine organisms based on the MASST search, the biochemical function of these molecules is of particular interest. The role of acylcarnitines in cellular energetics is well established; the varied detection of acylcarnitines in corals may indicate variability among species in their ability to readily utilize these pathways. Future work will focus on structural description of these carnitines. How the profiles of metabolites reported in this manuscript change over time should also be characterized to determine their viability as biomarkers of health, disease, and lesion progression. The juxtaposition of *M. meandrites* SCTLD susceptibility and the observed highest bioactivity of the BuOH partitions of the extracts of this species against putative secondary SCTLD pathogens generates interesting avenues for future study, including how molecular dynamics of pathogen response and disease susceptibility might explain discrepancies between disease susceptibilities in the field while metabolite extracts show high antibacterial activity when challenged with in lab-assays to a specific pathogen. Additional studies can also incorporate knowledge of environmental factors like heat stress to determine how biochemical disease dynamics and susceptibility may shift in the field.

As SCTLD appears to affect Symbiodiniaceae and disrupt their relationship with the host(13, 38, 40, 43), it is imperative to understand the differences in chemical cross-talk between the corals and endosymbionts. Symbiodiniaceae dynamics within the host (e.g., relative abundance, density, species) will likely have an effect on the metabolomic profiles. In this study, several Symbiodiniaceae metabolites driving interspecies differences and endosymbiont-derived carotenoid pigments displayed both inter- and intraspecies variation suggesting the presence of different endosymbiont genotypes in these samples and/or different endosymbiont-host dynamics among species. Thus, metabolomic studies on Symbiodiniaceae directly isolated from field corals will enable source tracking to tease apart host-derived, diet-derived and endosymbiont-derived compounds. Given that the endosymbiont fraction can be isolated from corals using mechanical methods such as the air brush technique (124), we propose future studies also include comprehensive metabolic profiling of these endosymbionts prior to and upon exposure to disease. Such differences should be interrogated across corals with different disease susceptibility and with different endosymbiont profiles. Furthermore, studies of metabolomic profiles taken at discrete time-points after disease exposure will provide insights into the transitory response of corals to disease stressors. With the increased application of untargeted metabolomics methods to study coral physiology, availability of annotated datasets, and our ability to mine public datasets using methods such as MASST, discoveries pertaining to chemical interactions between the coral host, the endosymbiont, environment, and the microbiome that define health status are a real possibility. By advancing our knowledge of the biochemical pathways involved in coral health and susceptibility to disease, we can disentangle the sources of metabolites, and how they change with time and increasing anthropogenic and climate related stressors.

## Materials and Methods

### Coral Sample Collection and Extraction Procedure

Whole coral fragments of a maximum size of 200 cm^2^ were collected on SCUBA from four visually healthy Scleractinia coral species, *O. faveolata* (n=4)*, M. cavernosa* (n=4)*, M. meandrites* (n=4), and *C. natans* (n=3). These were collected in January 2020 from four sites outside of the Dry Tortugas National Park (Figure 1 and Table S1). This collection occurred ahead of the SCTLD front, which was first observed at the Dry Tortugas in May 2021(125). Collection also occurred during a time of year when temperature stress and any associated paling or bleaching of the corals should not have been occurring, and none was observed at time of collection. Corals were chiseled at the base until they released from the substrate and then were transported back to the diving vessel in 18.9 L plastic bags filled with ocean water. Collected corals larger than 25 cm^2^ were cut down to this size on the diving vessel using an AquaSaw (Gryphon C-40 CR). The cut portions and whole colonies were stored in a 1000 L covered insulated container (Bonar Plastics, PB2145) filled halfway with ocean water. Air stones within the container allowed for aeration and water movement. A complete water change was performed on the container four times daily. Collections were conducted over two days before corals were transported the morning of the third day. The cruise was sponsored by the Florida Department of Environmental Protection and sample collection was covered by permit FKNMS-2019-160 to Valerie Paul. All corals collected from the field were transported to the Smithsonian Marine Station in Fort Pierce, FL. For transport, individual colonies were wrapped in plastic bubble wrap moistened with ocean water and then placed in a cooler.

Upon arrival, corals were rinsed with filtered seawater (FSW) and stored in a large indoor recirculating system holding approximately 570 L of FSW at 25.5 °C ± 0.3 °C. The FSW was collected from an intake pipe extending 1600 m offshore South Hutchingson Island, Port Saint Lucie, FL and was filtered progressively through 20, 1.0, 0.5, and 0.35 µm pore filters. While stored prior to use in the recirculating system, the FSW constantly circulated through a 20 µm pore filter, a filter canister with ROX 0.8 aquarium carbon (Bulk Reef Supply), and a 36-watt Turbo-twist 12x UV sterilizer (Coralife) in series. The recirculating system contained a UV sterilizer (same model as described), two circulating pumps (AquaTop MaxFlow MCP-5) to create water movement, and a row of 6 blue-white 30 cm^2^ LED panels (HQPR) to create 150 to 250 µmol photons m^-2^ sec^-1^ for the captive corals. Corals were stored in the recirculating system for 5 days prior to sampling to allow them time to recover after transport, with a partial water change on the fourth day. All corals were held together in a single table.

After the fifth day, the corals were cut into smaller segments with a rock saw, and the blade was constantly sprayed down with UV/filter-sterilized seawater to cool the blade and wash off any debris, thus reducing cross-contamination between corals. The coral fragments ranged from 1-13 cm^2^ in surface area (6.2+3.7 cm^2^, mean+SD) were immediately frozen (-80 °C) and lyophilized the next day. Coral fragments were lyophilized overnight and then extracted twice using a 2:2:1 mixture of ethyl acetate (EtOAc), methanol (MeOH) and water (H_2_O) at room temperature in 20 mL scintillation vials. For the extraction, coral fragments were covered with an excess of solvent mixture, sonicated for 5-10 min and left to sit for 3 h on the initial extraction and overnight for the second extraction. The liquid extract was then transferred into a round-bottom flask using filter paper to prevent the transfer of coral fragments. The coral extract was then dried via rotary-evaporation (Buchi Rotovapor R-210) in a 35 °C water bath (Buchi Heating Bath B-491) and weighed to determine the amount of crude extract. The extracts were dried *in vacuo* and 0.5 mg of the extract was transferred to Eppendorf tubes for metabolomics data analysis. The extracts were stored at −20 °C until UPLC-MS data was acquired.

### Endosymbiont Metabolome Data

We previously reported on metabolome profiles of Symbiodiniaceae isolates provided by Mary Alice Coffroth from the University of Buffalo Undersea Reef Research (BURR) collection (43). Given the challenges involved in isolating and culturing Symbiodiniaceae, we used this publicly available data(43) (massIVE identifier MSV000087471) in this current study to aid in determining the biosynthetic producer of detected metabolites. Briefly, the endosymbionts were isolated by Mary Alice Coffroth from Orbicella faveolata corals sampled between 2002-2005 from the Florida Keys. Isolate extracts were sent by Richard Karp and Andrew Baker, (University of Miami). Culture conditions included incubation at 27°C in F/2 media, with 20 μE of light on a 14:10 diurnal cycle. Extracts of the culture were performed as previously described(43), using 2:2:1 EtOAc:MeOH:H_2_O to extract pelletized cellular cultures. Solvents were removed and the dried samples were transferred in 3:1 MeOH:H_2_O into a 1.5 mL microcentrifuge tube. After centrifugation, the supernatant was transferred to a microcentrifuge tube and removed via SpeedVac for 3 h. The extract was frozen, lyophilized, and sent to the Georgia Institute of Technology for metabolomics data acquisition and analysis. Please see Deutsch *et al* 2021 for complete methodology(43).

### Mass Spectrometry Data Acquisition and Analysis

The dried extracts were resuspended in 100% MeOH containing 1 μM sulfadimethoxine as an internal standard. The samples were analyzed with an Agilent 1290 Infinity II UHPLC system (Agilent Technologies) using a Kinetex 1.7 μm C18 reversed phase UHPLC column (50 × 2.1 mm) for chromatographic separation, coupled to an ImpactII ultrahigh resolution Qq-TOF mass spectrometer (Bruker Daltonics, GmbH, Bremen, Germany) equipped with an ESI source for MS/MS analysis. MS/MS spectra were acquired in positive mode as previously described(43). Metabolomics data on the cultured Symbiodiniaceae from the Burr Collection was previously acquired(43). The strains utilized are reported in Table S5.

The raw data was converted to mzXML format using vendor software. MZmine 2.53 was used to extract metabolite features with steps for mass detection, chromatogram building, chromatogram deconvolution, isotopic grouping, retention time alignment, duplicate removal, and missing peak filling(126). This processed data was submitted to the feature-based molecular networking workflow on the Global Natural Product Social Molecular Networking (GNPS) platform(127). The output of MZmine includes information about LC-MS features across all samples containing the *m/z* value, retention time, the area under the peak for the corresponding chromatogram, and a unique identifier for each feature. The quantification table and the linked MS² spectra were exported using the GNPS export module(126, 128) and the SIRIUS 4.0 export module(126, 129). Feature Based Molecular Networking was performed using the MS² spectra (.mgf file) and the quantification table (.csv file). The molecular network was generated as previously described(43). The molecular network and the generation parameters are available <u>here</u>. The molecular network was visualized using Cytopscape v3.7.2(130). The MS2LDA analysis was performed as previously described with default parameters(131, 132). The MassQL Sandbox Dashboard(133) (v 0.3) on the GNPS platform was used to construct the spectral pattern queries for the MassQL search. Feature annotation was performed using SIRIUS with CSI:FingerID, MolDiscovery(134), GNPS spectral library matching, MassQL, MASST, and literature searches. The metabolite annotations presented herein follow the “level 2” annotation standard based upon spectral similarity with public spectral libraries, spectra published in the literature, and through spectral comparison with the analytical standards as proposed by the Metabolomics Society Standard Initiative(135). All mzXML files included in this study can be accessed publicly on the repository Mass Spectrometry Interactive Virtual Environment (MassIVE) with ID MSV000089633. The commercial analytical standard for α-tocopherolquinone (catalog number 35365) was purchased from Cayman Chemical Company and the commercial analytical standard for fucoxanthin (catalog number 16337) was purchased from Sigma Aldrich.

Prior to statistical analysis, blank subtraction was performed as previously described(43) to filter out features detected in the solvent and media controls. Unsupervised multivariate statistical analyses including principal component analysis(136) and hierarchical clustering analysis(137) were performed using MetaboAnalyst 5.0(138) and pareto scaling was employed prior to the analyses. The Plotter Dashboard (v.0.5) on the GNPS platform was used to construct boxplots for metabolite features of interest. A nonparametric Kruskal Wallis test with Dunn’s posttest was performed in R studio. The UpSet Plots were made using the Intervene app(139).

To test for an effect of coral species on metabolomic variation, we used a permutational multivariate analysis of variance (PERMANOVA)(62). Coral species was treated as a fixed effect (four levels), with site included as a random nested effect (four levels). The PERMANOVA was based on a Bray-Curtis similarity matrix(140), type III (partial) sums of squares, and 999 random permutations of square-root transformed data (to down-weight heavily dominant variables) under a reduced model. Both PERMANOVA and non-metric multidimensional scaling(141) plot were constructed from a Bray-Curtis similarity matrix of square root transformed data, which was performed using PRIMER v7(142). To quantify metabolomic variation within and between coral species, we calculated their multivariate dispersion using the PERMDISP routine (143). PERMDISP calculates the distance of each observation (in this case each coral sample) to its group centroid (in this case each coral species) and then compares the average of these distances among groups. It is a multivariate extension of Levene’s test, with the p-values obtained using permutations of the raw data. This allowed us to make inferences about the relative size of the clouds in multivariate space within and between coral species. The tests were based on the same transformed data and Bray-Curtis similarity matrix as our PERMANOVA tests. Shannon entropy(144) was calculated for the alpha diversity metric using a Jupyter Notebook. A Bray-Curtis similarity matrix of log transformed data was constructed using Primer v7 for the beta diversity metric. A principal coordinates analysis(145) was constructed on the Bray-Curtis similarity matrix. Several in silico tools were used to aid in metabolite annotations and are described herein. MolDiscovery compares in silico generated MS^2^ spectra of small molecules to user-uploaded experimental MS^2^ spectra(134). SIRIUS computes putative chemical formulas based on user-uploaded MS^1^ isotopic peaks and MS^2^ fragmentation patterns(129). CSI:FingerID transforms MS^2^ spectra into predicted structural fingerprints that enable matching to fingerprints generated from structure databases(146). CANOPUS, which predicts the chemical class of metabolites by utilizing CSI:FingerID’s predicted structural fingerprints, proposed the chemical class of 449.398_21.3 min as Vitamin E compounds(147). Unsupervised substructure discovery performed through MS2LDA(75) enabled annotations of several classes. Substructure motif 108 consisted of fragmentation peaks characteristic of tocopherol substructure (Figure S3D). Motif 185 containing characteristic carnitine headgroup fragment peaks(91, 148) at *m/z* 85.028 and 144.102 (Figure 4D) aided acylcarnitine annotations. Supervised substructure discovery was performed using MassQL, a MS query language platform that outputs metabolite features based on sets of user-defined fragment peaks and neutral losses(133).

### Extract Partitioning of Crude Extracts of Coral Metabolomes

Crude coral extracts were partitioned to remove salts and separate compounds based on polarity. First, approximately 3 mL of EtOAc was added to 20 mL scintillation vials containing dry crude coral extracts. Vials were sonicated to disrupt dry extracts for 30-60 seconds. 3 mL of H_2_O and another 1 mL of EtOAc was then added and the vials swirled to mix. Vials were then left to separate into distinct layers. The EtOAc layer was then transferred via glass pipette into a clean and pre-massed 20 mL scintillation vial. An additional 2 mL of EtOAC was then added into the crude mixture with water, swirled to mix and left to separate again. The EtOAc layer was again transferred into the vial containing the EtOAc partition. The EtOAc partitions were then dried via a SpeedVac vacuum concentrator (Thermo Scientific Savant SPD121P) at 35 °C. The remaining coral extract water mixture was then partitioned using n-butanol (BuOH). Approximately 2 mL of BuOH was added to the water extract mixture, swirled to mix and then left to sit until distinct layers formed. The BuOH partition was then transferred into a clean and pre-massed 20 mL scintillation vial. Another round of BuOH partitioning was then done by adding an additional 1 mL of BuOH to the water mixture, mixing and allowing time for a final separation. After the BuOH layer was transferred to the vial containing the initial BuOH partition, it was dried via rotary-evaporation and sent for LC-HRMS data acquisition. The BuOH partitions were resuspended and LC-MS data was acquired and analyzed following the procedure outlined in “Mass Spectrometry Data Acquisition and Analysis”.

### Disk Diffusion

Coral extracts were tested for antibacterial activity using disk diffusion growth inhibition assays against putative coral pathogens, *Vibrio coralliilyticus* OfT6-21 and *V. coralliilyticus* OfT7-21, *Leisingera* sp. McT4−56 and *Alteromonas* sp. McT4-15. To make pathogen lawns, overnight liquid cultures of pathogens were grown by inoculating 2-3 mL of seawater broth (4 g/L tryptone and 2 g/L yeast extract in 0.22 mm filtered seawater) with individual colonies of each strain and shaking culture tubes at 150 RPM and 28 °C (Benchmark Incu-shaker 10LR). To coat seawater agar (seawater broth with 15 g/L agar) plates with a pathogen lawn, a 200 mL aliquot of liquid culture (OD600 =0.5) was added to each plate (150 mm x 15 mm) and spread using sterile glass beads.

Coral partitions were tested by solubilizing partitions in MeOH to a concentration of 6.25 mg/mL and applying 4 µL aliquots to sterile paper disks (Whatman Grade 1) in triplicate (final amount 25 µg/disk). A filter disc with 4 μL of nalidixic acid at 15.62 mg/mL (62.5 μg) was used as a positive control. Negative controls were disks treated with MeOH only. Disks were given time to dry completely and then carefully transferred with sterile forceps to seawater agar plates containing freshly coated pathogen lawns. Disk diffusion plates were then incubated at 28 °C overnight. After incubation, zones of inhibition (ZOI) were measured using digital calipers from the edge of the paper disk to the edge of the zone of bacterial growth inhibition.

### Ethics

The sample collection was covered by permit FKNMS-2019-160 to Valerie Paul.

## Supporting information

All supplemental files

## Data accessibility

The metabolomics data utilized in this manuscript is available at gnps.ucsd.edu with MassIVE ID# MSV000089633. The data acquired in negative mode is also available in this dataset, but is not presented in this study due to the lack of high-confidence annotations. The code utilized in this manuscript is available at https://github.com/Garg-Lab/Dry-Tortugas-Corals-Files.

## Acknowledgements

We thank Olivia Carmack for assistance collecting the corals in the Dry Tortugas and Woody Lee, Blake Ushijima and Jay Houk for assistance cutting and caring for the corals in aquaria. We thank Monica Monge Loria for providing an image of cultured Symbiodiniaceae used in Figure 1. We also thank the multi-agency effort funded by the Florida Department of Environmental Protection to collect corals. Healthy corals were collected for this study under Florida Keys National Marine Sanctuary Permit Number FKNMS-2019-160.

