## Supplementary material for "Metabolomic profiles of stony coral species from the Dry Tortugas National Park display inter- and intraspecies variation": All supplemental files

**Running title:** Intra- and interspecies variation in coral metabolomes

### Table of Contents


Figure S1. Unsupervised multivariate statistical analyses of coral extracts .............................. **S3**

Figure S2. Alpha and beta diversity metrics ............................................................................ **S4**

Figure S3. Non-metric multi-dimensional scaling analysis ....................................................... **S5**

Figure S4. $MS^2$ spectra analyses of α-tocopherolhydroquinone and α-tocopherolquinone ....... **S6**

Figure S5. Acylcarnitine annotation and differential detection ................................................ **S7**

Figure S6. Annotation of DGCC betaine lipids ......................................................................... **S8**

Figure S7. $MS^2$ spectral analysis of lyso-DGCC(16:0) and analogues.................................... **S9**

Figure S8. $MS^2$ mirror plots of features annotated as pigments ............................................ **S10**

Figure S9. Bioactivity of BuOH partitions and annotations of compounds detected................ **S11**

Figure S10. $MS^2$ Mirror Plots for annotations of compounds uniquely detected in BuOH partitions ................................................................................................................................ **S12**

Links to MassQL Query Output ............................................................................................... **S12**

Table S1. . ................................................................................................................................ **S13**

Table S2. ............................................................................................................................. **S14-15**

Table S3. ................................................................................................................................. **S16**

Table S4. ............................................................................................................................ **S17-S30**

Table S5. ................................................................................................................................. **S30**

References. .............................................................................................................................. **S30**

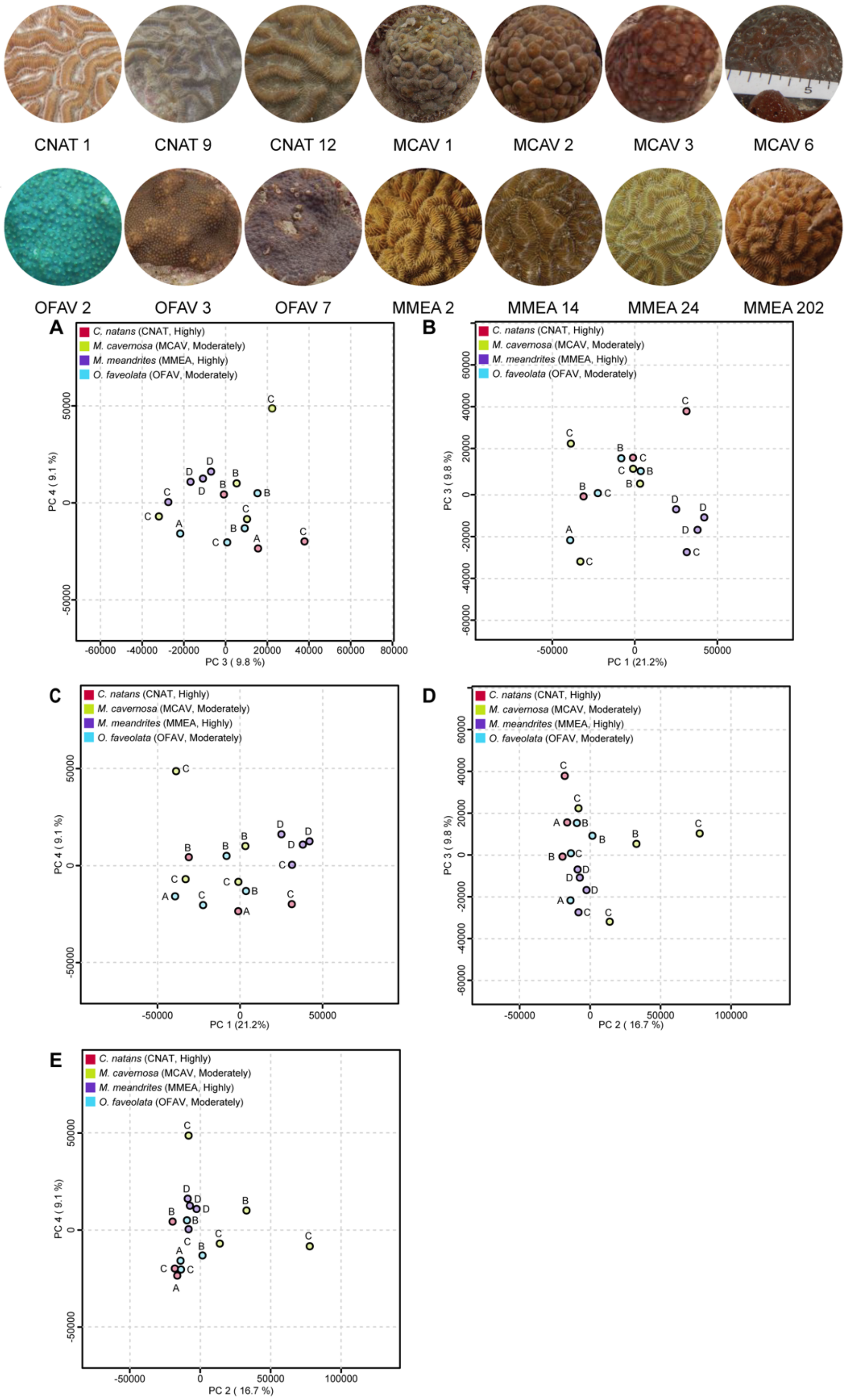


**Figure S1**. **Images of coral colonies collected and analyzed in this study (top) and unsupervised multivariate statistical analyses of coral extracts**. **(bottom A-E)** The first four principal components from the PCA model. The variation explained by the principal components is included on the corresponding axes. The metabolome profiles of *M. meandrites* show tight clustering across all components, implying low intraspecies variation. For the other species, the principal components shown capture intraspecies variation. The SCTLD susceptibility categorization is included in the key ('Highly', 'Moderately').

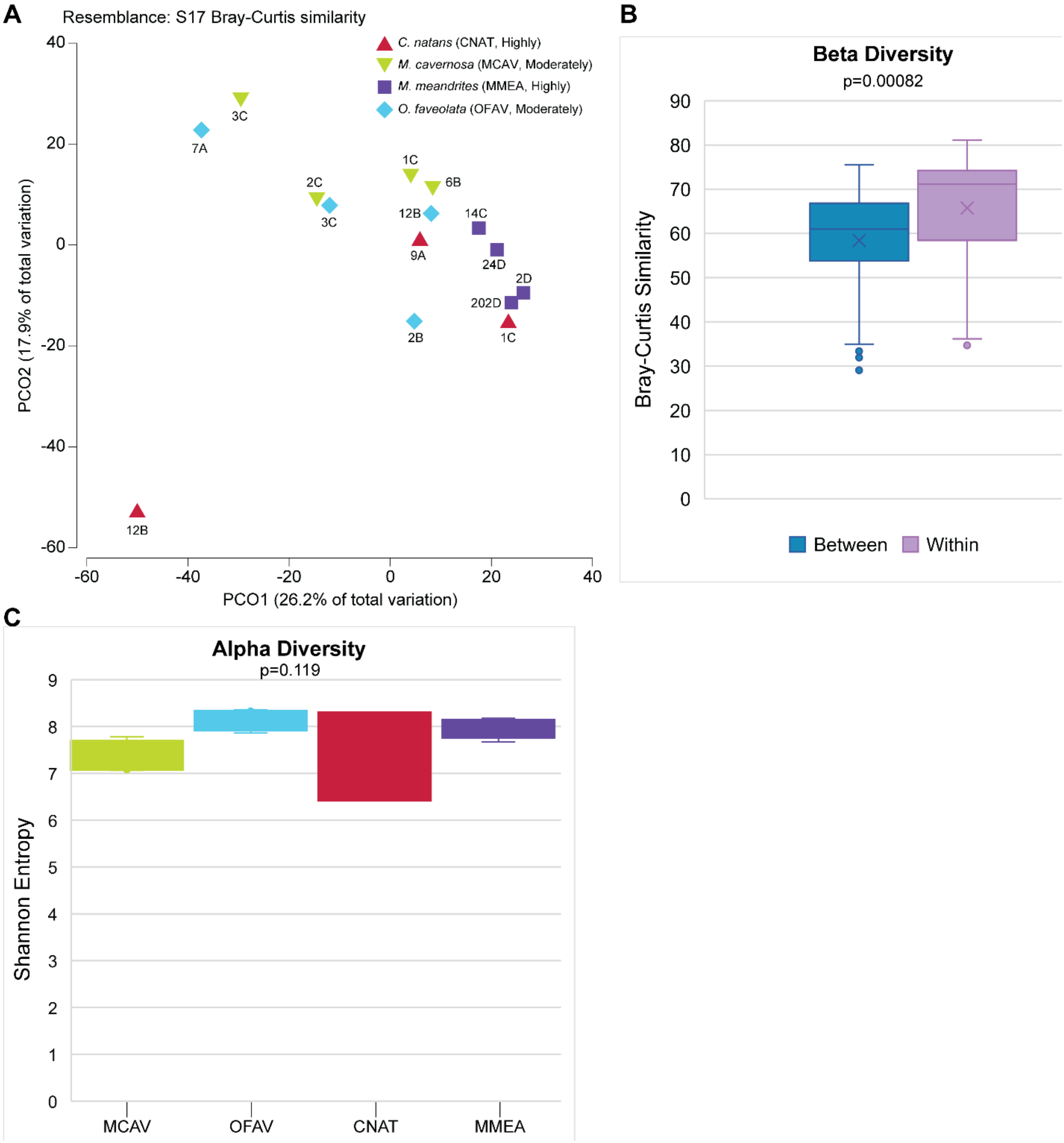


**Figure S2. Alpha and beta diversity metrics. (A)** Principal coordinates analysis constructed on the Bray-Curtis similarity matrix of the metabolome data. **(B)** Plot of the Bray-Curtis similarity scores between and within species. The Bray-Curtis similarity scores within species is significantly larger (p=0.00082) than the scores between species. **(C)** Plot of the alpha diversity metrics by coral species. No significant comparison was found using a Kruskal-Wallis test. The tests were conducted on a small sample size.

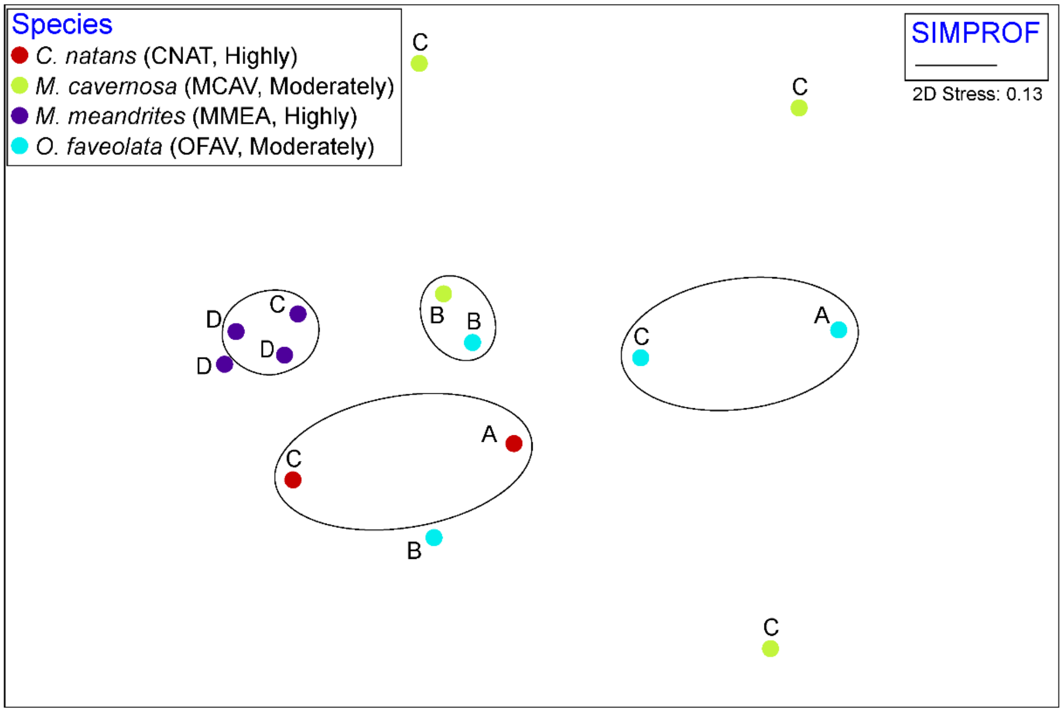


**Figure S3**. **Non-metric multi-dimensional scaling analysis.** n-MDS plot shows clustering of data into four groups with one group consisting of all *M. meandrites* samples. The SCTLD susceptibility categorization is included in the key (‘Highly’, ‘Moderately’). The largest variation is observed for *M. cavernosa*. Initial analysis using SIMPROF cluster and nMDS revealed CNAT12 as an outlier; thus it was removed from this analysis. Permutational analysis of variance found significance between species (p=0.005) but not sample site.

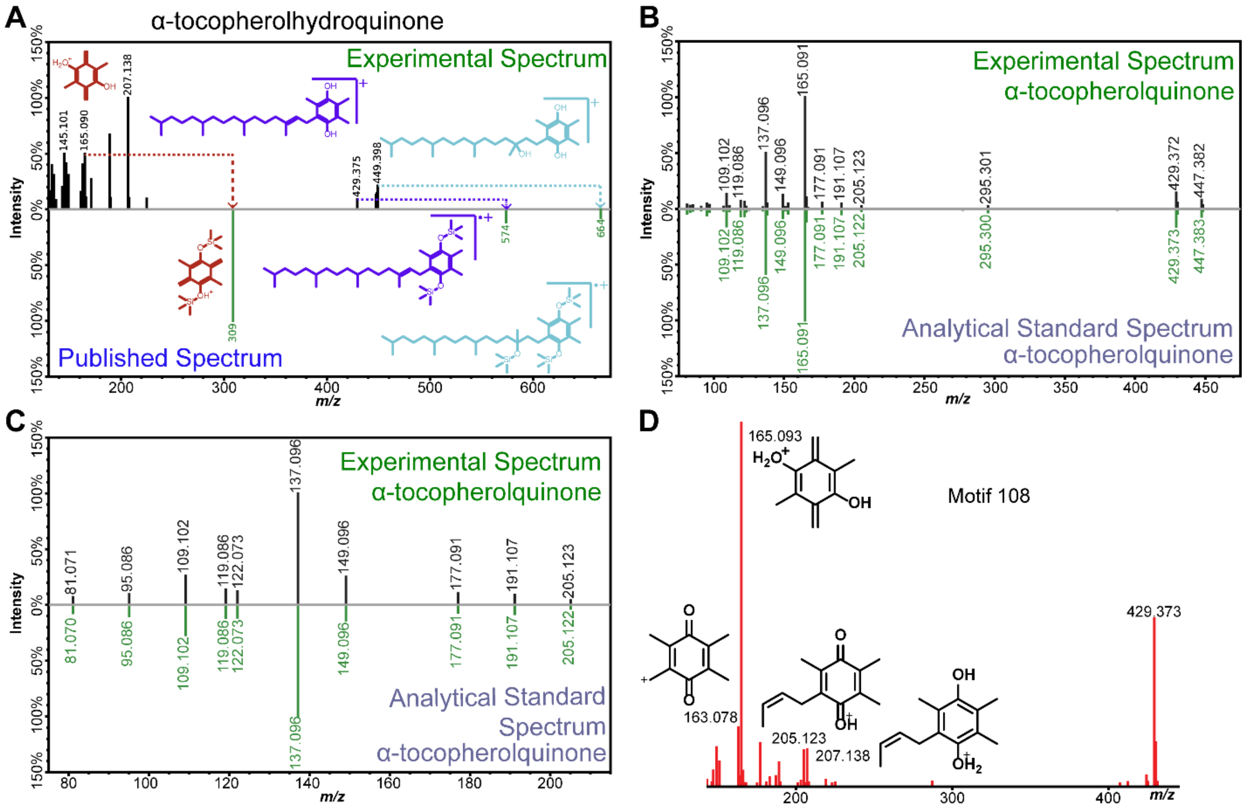


**Figure S4. MS² spectra analyses of α-tocopherolhydroquinone and α-tocopherolquinone. (A)** MS² mirror plot comparing the experimental spectrum of α-tocopherolquinone (top) compared to a published(1) spectrum of silicated α-tocopherolquinone (bottom). Key chemical substructures supporting the annotation are included, where substructures of the same color correspond. **(B)** MS² mirror plot comparing the experimental spectrum of *m/z*_RT 447.383_21.3 (top) with the spectrum acquired on an analytical standard of α-tocopherolquinone (bottom). **(C)** MS² mirror plot of (B), zoomed in to the *m/z* 80-215 range where the fragment peak at *m/z* 165 has been removed to show matching fragment peaks detected at lower intensity. **(D)** MS2LDA motif 108 used to aid annotations. Chemical substructures for characteristic fragment peaks are included.

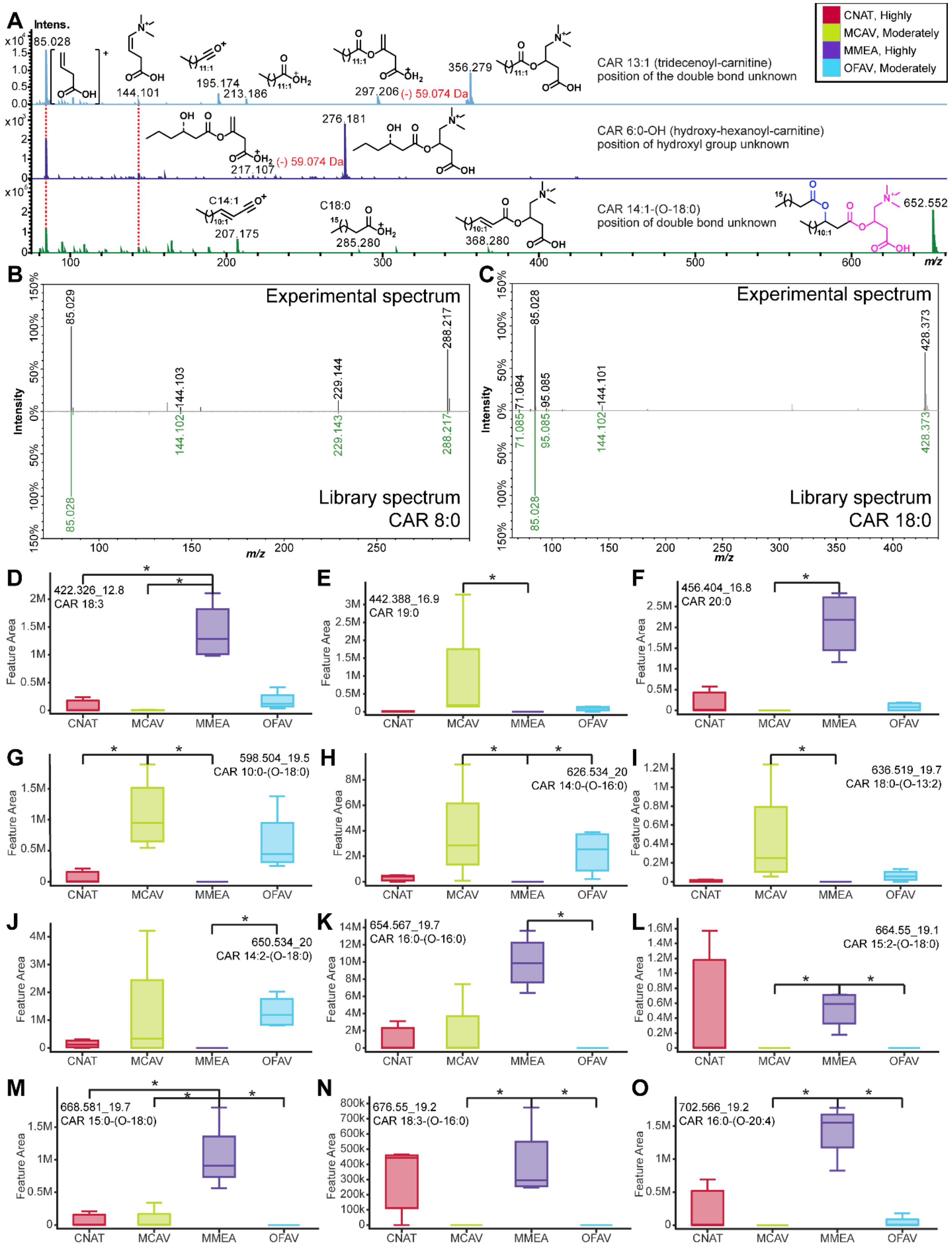

A
B
C
D
E
F
G
H
I
J
K
L
M
N
O
CNAT, Highly
MCAV, Moderately
MMEA, Highly
OFAV, Moderately
CAR 13:1 (tridecenoyl-carnitine)
position of the double bond unknown
CAR 6:0-OH (hydroxy-hexanoyl-carnitine)
position of hydroxyl group unknown
CAR 14:1-(O-18:0)
position of double bond unknown
(-) 59.074 Da
Experimental spectrum
Library spectrum
CAR 8:0
CAR 18:0
422.326_12.8
CAR 18:3
442.388_16.9
CAR 19:0
456.404_16.8
CAR 20:0
598.504_19.5
CAR 10:0-(O-18:0)
626.534_20
CAR 14:0-(O-16:0)
636.519_19.7
CAR 18:0-(O-13:2)
650.534_20
CAR 14:2-(O-18:0)
654.567_19.7
CAR 16:0-(O-16:0)
664.55_19.1
CAR 15:2-(O-18:0)
668.581_19.7
CAR 15:0-(O-18:0)
676.55_19.2
CAR 18:3-(O-16:0)
702.566_19.2
CAR 16:0-(O-20:4)
Feature Area
CNAT
MCAV
MMEA
OFAV

**Figure S5**. **Acylcarnitine annotation and differential detection. (A)** The $MS^2$ spectra of representative acylcarnitine analogues are shown. Conserved fragment peaks, with the chemical substructure, are indicated between spectra. Key fragment peaks used to support the acylcarnitine annotation include *m/z* 85.028 and 144.101.(2) A representative $MS^2$ spectra of an acylcarnitine analogue containing a conjugated fatty acyl ester of hydroxy fatty acid (FAHFA) is also shown (bottom spectrum) The acylcarnitine headgroup is shown in pink, and the novel linkage of the fatty acyl ester is in blue. The fragment observed at *m/z* 285.280 is representative of C18:0 and supports this annotation. **(B)** and **(C)** $MS^2$ mirror plots for the experimental spectrum (top) and the library spectrum in GNPS (bottom). **(D-O)** The box plots of the relative abundance of features annotated as acylcarnitines. Asterisks indicate significant differences between the compared groups as determined by a Kruskal Wallis test with Dunn's post-hoc test (adjusted $p<0.05$). Each box plot is labeled with the *m/z*_RT and proposed annotation. The SCTLD susceptibility categorization is included in the key at the top of the figure ('Highly', 'Moderately').

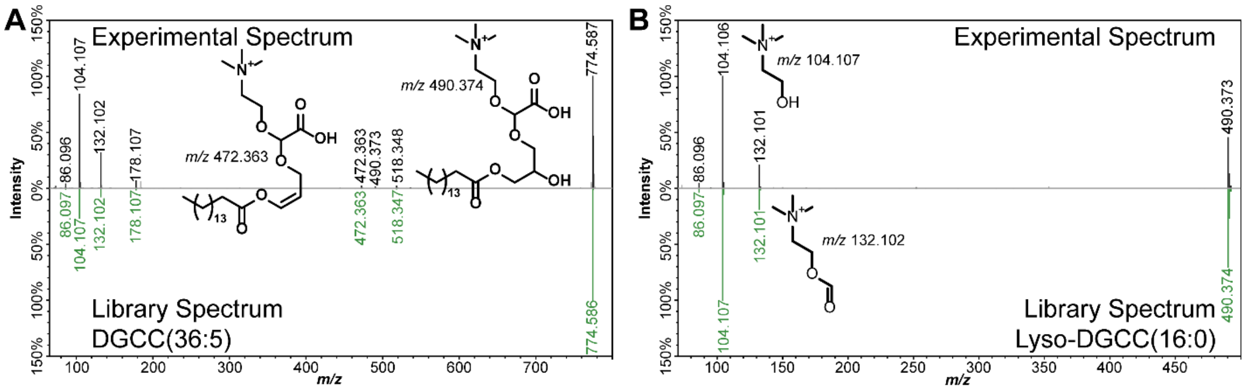


**C**

```
QUERY scaninfo(MS2DATA) WHERE
MS2PROD=104.1070:TOLERANCEPPM=5
AND MS2PROD=132.1019:TOLERANCEPPM=5
AND MS2PROD= 472.3633:TOLERANCEPPM=5
AND MS2PROD=490.3738:TOLERANCEPPM=5
```

**Figure S6. Annotation of DGCC betaine lipids. (A)** and **(B)** $MS^2$ mirror plots for the experimental spectra (top) and the library spectra in GNPS (bottom). Key fragment peaks at *m/z* 490.373 and 472.373 enabled (A) to be further annotated as DGCC(16:0_20:5) (Figure S6). The corresponding chemical substructures for these fragment peaks are included, including substructures for *m/z* 132.102 and 104.107. **(C)** Shows the query submitted to MassQL to search for lyso-DGCC(16:0) analogues containing the 16:0 fatty acid tail.

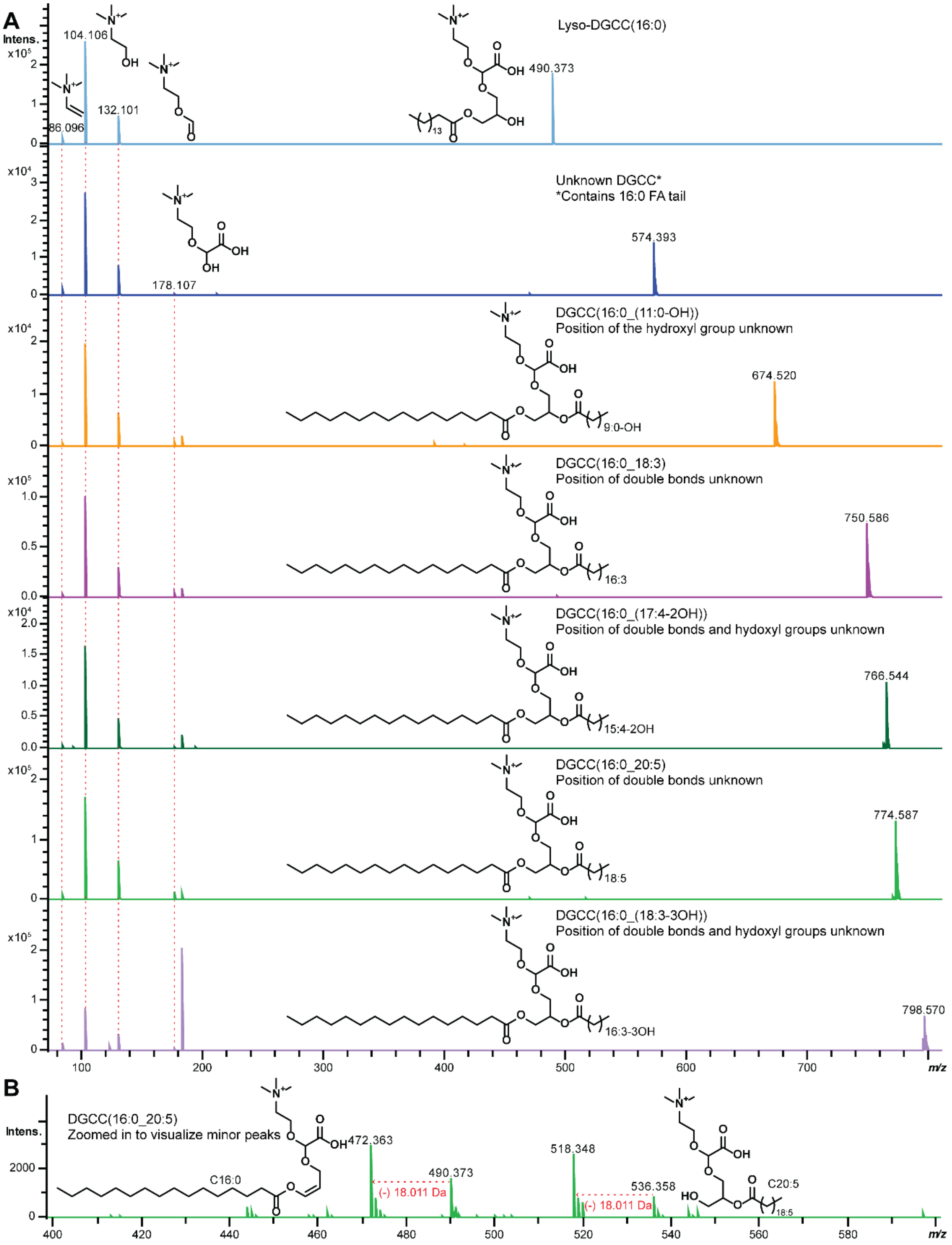


**Figure S7. MS² spectral analysis of lyso-DGCC(16:0) and analogues. (A)** The MS² spectra of lyso-DGCC (monoacylated; 16:0) and diacylated DGCCs are shown. Conserved fragment peaks, including the chemical substructure, are indicated between spectra. Fragment peaks *m/z* 86.096, 104.107, 132.102, and 178.107 support the DGCC annotation. Mass shifts between key fragment peaks that aided in annotation are indicated. **(B)** A zoomed in of the spectrum of DGCC (16:0_20:5) in the *m/z* range 400-600 is shown. The fragment peaks at *m/z* 490.373 and 472.363 support the 16:0 fatty acyl tail.

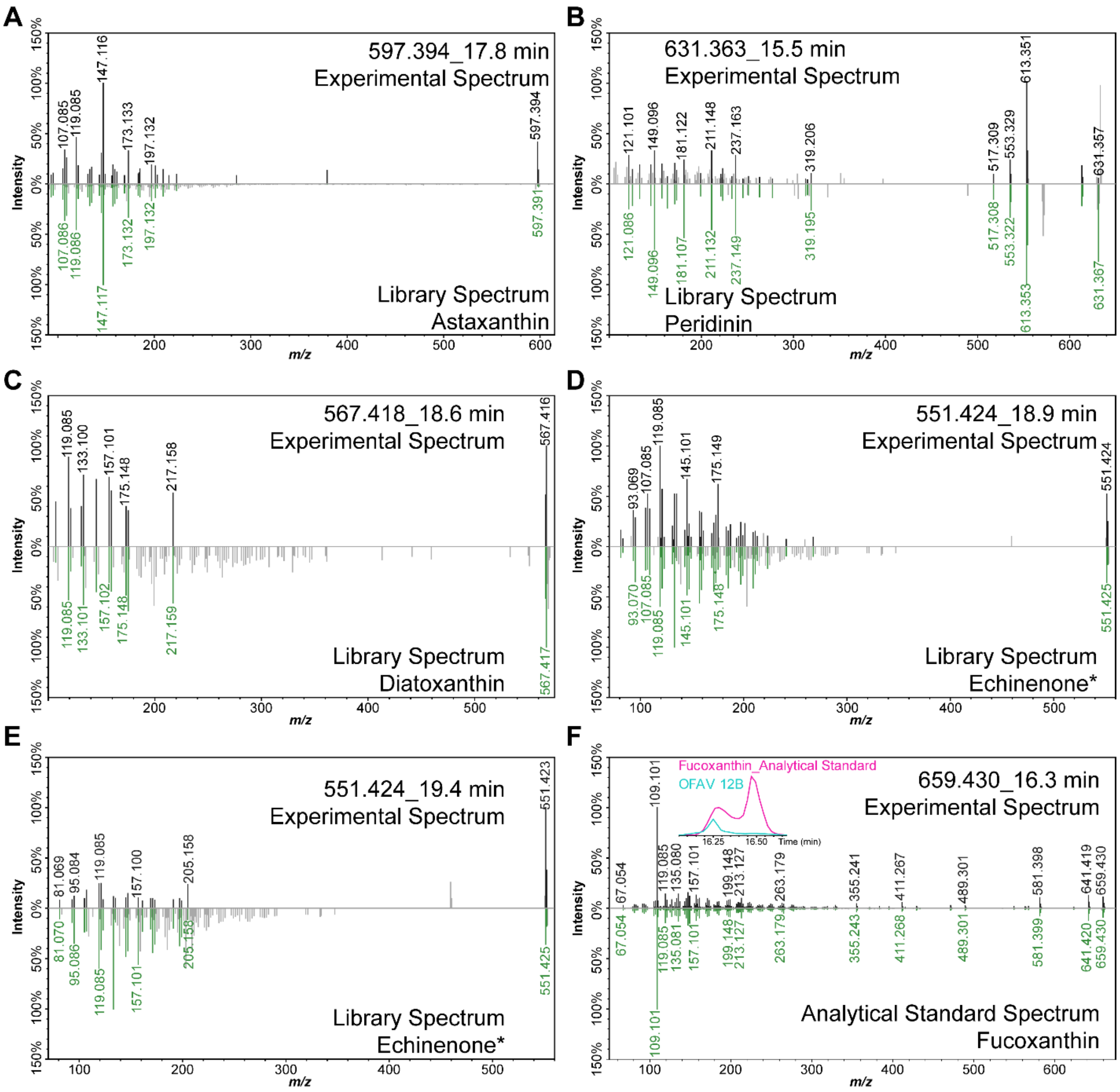


G

| *m/z*_RT | Putative Annotation | *Breviolum* | *Durusdinium* | *Symbiodinium* |
|---|---|---|---|---|
| 551.424_19.4 | Echinenone or isomer | A | A | A |
| 551.425_18.9 | Echinenone or isomer | A | A | A |
| 567.418_18.6 | Diatoxanthin | A | A | A |
| 583.415_18.1 | Diadinoxanthin or Diadinochrome* | A | A | A |
| 583.415_20.5 | Diadinoxanthin or Diadinochrome* | P | P | A |
| 585.43_17 | Antheraxanthin | A | A | A |
| 597.394_17.8 | Astaxanthin | A | A | A |
| 601.423_15.1 | Neoxanthin | A | A | A |
| 613.352_17.7 | Pyrroxanthin | P | P | A |
| 631.363_15.5 | Peridinin | P | P | P |
| 659.430_16.3 | Fucoxanthin | A | P | A |
| 583.416_18.4 | Diadinoxanthin or Diadinochrome* | P | A | P |
| | | P: present | A: absent | |

**Figure S8. MS² mirror plots of features annotated as pigments. (A-E)** MS² mirror plots for the experimental spectra (top) and the library spectra in GNPS (bottom) are shown, along with the pigment name, *m/z* and retention time. The * indicates features that have identical MS² spectra, but different retention times representing isomeric species. **(F)** $MS^2$ mirror plot for the experimental spectrum of the feature annotated as fucoxanthin compared with the spectrum acquired on a fucoxanthin analytical standard. The extracted ion chromatogram for *m/z* 659.430 is shown as an inset for an *O. faveolata* extract and the analytical standard. **(G)** The list of annotated pigments, including their detection pattern in the zooxanthellae genera included in this study.

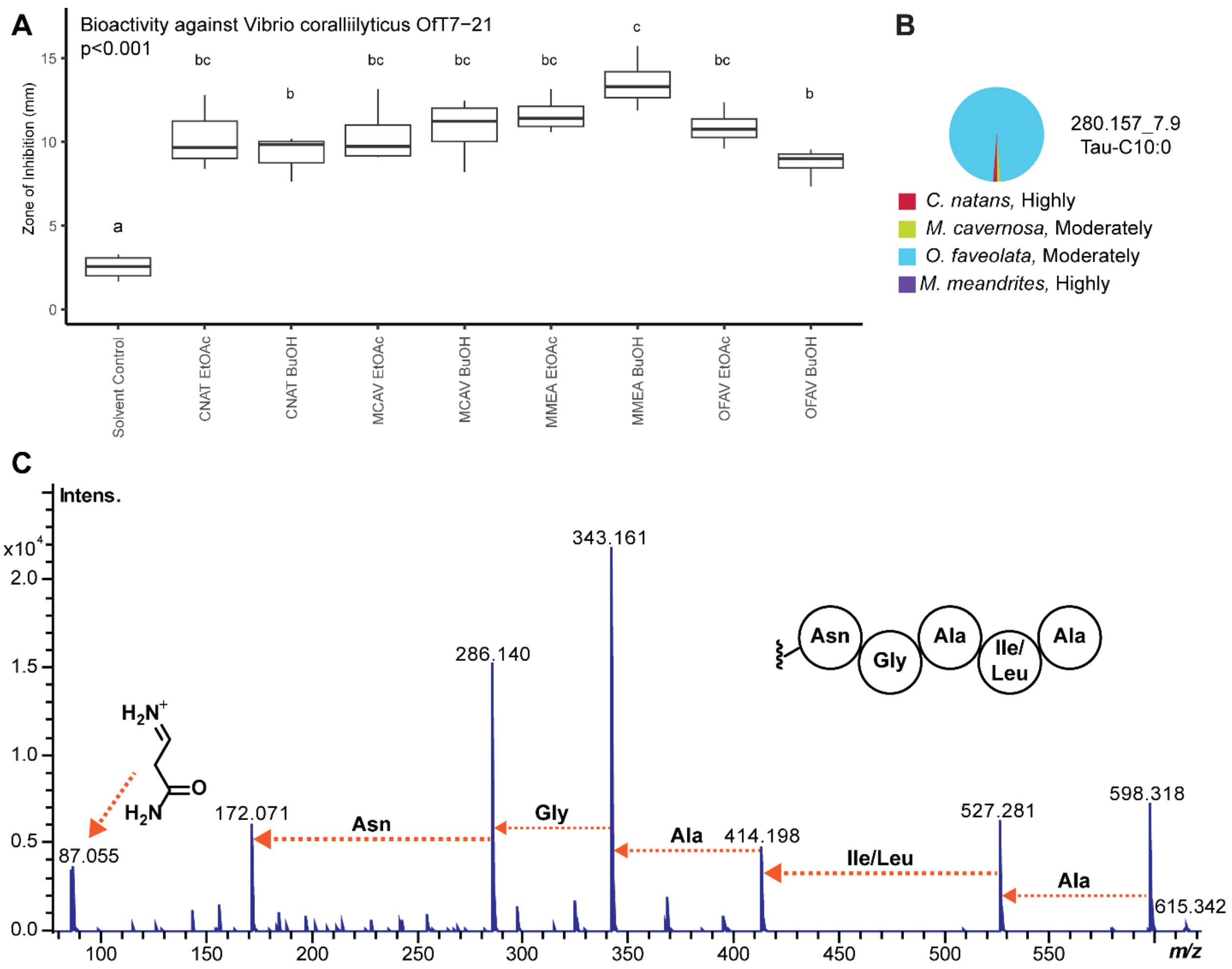


**Figure S9. Bioactivity of BuOH partitions and annotations of compounds detected. (A)** Bioactivity of EtOAc and BuOH partitions of crude extracts determined using agar diffusion growth inhibition assay against the coral pathogen *V. coralliilyticus* OfT7-21. **(B)** The feature annotated as Tau-C10:0 showed highest detected relative abundance in *O. faveolata*. The SCTLD susceptibility categorization is included in the key (‘Highly’, ‘Moderately’). **(C)** Partial annotation of a polypeptide. Feature 615.3461_5.9 min was proposed as an analogue of tunicyclin G by DEREPLICATOR. The experimental $MS^2$ spectrum supports the annotation of a polypeptide containing amino acid residues NGAI/LA. The detection of the Asn immonium ion (*m/z* 87.055) aided the annotation and refuted the annotation of a tunicyclin G analogue.

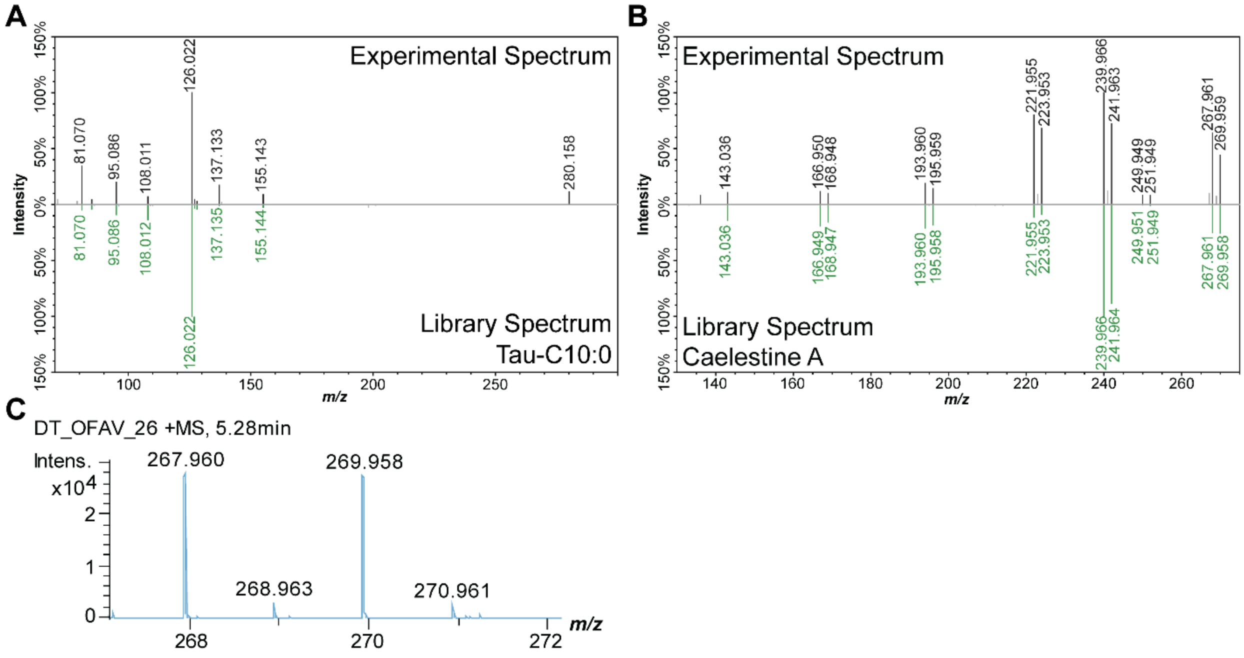


**Figure S10. MS$^2$ Mirror Plots for annotations of compounds uniquely detected in BuOH partitions. (A)** Tau-C10:0 was annotated through the GNPS library where the experimental spectrum (top) is compared to the GNPS library spectrum (bottom). **(B)** MS$^2$ mirror plot of the feature annotated as Caelestine A, where the experimental spectrum (top) is compared to the GNPS library spectrum (bottom). **(C)** Isotopic pattern representative of bromination.

**Links to MassQL Query Output:**

Acylcarnitine analogues:
https://gnps.ucsd.edu/ProteoSAFe/status.jsp?task=a1029fbb7b2b49e787febf1942032b2f

Lyso-DGCC(16:0) analogues:
https://gnps.ucsd.edu/ProteoSAFe/status.jsp?task=8737be5237d445889bba22013b2cfd70

**Table S1.** Sample and collection site information.

| Species | Identification Number & Site | Depth (m) | Fragment Surface Area ($cm^2$ or cmxcm) | Date Collected |
|---|---|---|---|---|
| *M. meandrites* | 14, C | 14.6 | 7.98 | 1/28/2020 |
| *M. meandrites* | 24, D | 16.8 | 4 | 1/28/2020 |
| *M. meandrites* | 2, D | 13.4 | 1 | 1/28/2020 |
| *M. meandrites* | 202, D | 13.7 | 5.75 | 1/28/2020 |
| *C. natans* | 1, C | 13.4 | 2.1 | 1/28/2020 |
| *C. natans* | 9, A | 18.3 | 13.6 | 1/29/2020 |
| *C. natans* | 12, B | 17.4 | 5.6 | 1/28/2020 |
| *M. cavernosa* | 2, C | 14.3 | 9 | 1/28/2020 |
| *M. cavernosa* | 3, C | 14.3 | 11.2 | 1/28/2020 |
| *M. cavernosa* | 1, C | 14.3 | 5.4 | 1/28/2020 |
| *M. cavernosa* | 6, B | 16.8 | 3.4 | 1/28/2020 |
| *O. faveolata* | 2, B | 15.8 | 5x4 | 1/28/2020 |
| *O. faveolata* | 3, C | 14 | 6x3 | 1/28/2020 |
| *O. faveolata* | 7, A | 18 | 5x4 | 1/29/2020 |
| *O. faveolata* | 12, B | 16.5 | 5x5 | 1/28/2020 |

| **Site** | A | B | C | D |
|---|---|---|---|---|
| **Coordinates** | 24°38.564 N, 82°58.161 W | 24°35.608 N, 82°58.862 W | 24°35.275 N, 82°58.767 W | 24°35.000 N, 82°58.236 W |
| **Approximate Distance Between Sites (km)** | A | B | C | D |
| B | 6 | | | |
| C | 6 | 1 | | |
| D | 7 | 2 | 1 | |

**Table S2.** Metabolite features annotated in this study.

| *m/z*_RT | Putative Annotation | Annotation Method |
|---|---|---|
| **Tocopherol Analogues** | | |
| 429.373_20.5 | α-tocomonoenol* | Previous study[1] |
| 429.373_20.7 | α-tocomonoenol* | Previous study[1] |
| 429.373_21.7 | α-tocomonoenol* | Previous study[1] |
| 429.373_22.3 | α-tocomonoenol* | Previous study[1] |
| 447.383_21.1 | α-Tocopherolquinone* | SIRIUS with CSI:Finger ID |
| 447.383_21.3 | α-Tocopherolquinone* | FBMN Propagation, Analytical Standard[3] |
| 449.398_21.3 | α-Tocopherolhydroquinone | MolDiscovery and SIRIUS with CSI:Finger ID |
| 441.337_19.0 | Hydroxy-α-tocotrienol | SIRIUS with CSI:Finger ID |
| 424.333_19.7 | α-tocotrienol | Previous study[1] |
| **Acylcarnitines** | | |
| 276.18_2.8 | CAR 6:0-OH | MS2LDA |
| 288.217_7.4 | CAR 8:0 | GNPS Library Match |
| 344.279_11.1 | CAR 12:0 | MS2LDA |
| 356.279_11.2 | CAR 13:1 | SIRIUS with CSI:Finger ID |
| 372.311_12.6 | CAR 14:0 | SIRIUS with CSI:Finger ID |
| 396.311_12.2 | CAR 16:2 | MS2LDA |
| 398.326_13.1 | CAR 16:1 | SIRIUS with CSI:Finger ID |
| 400.342_13.9 | CAR 16:0 | MassQL |
| 422.326_12.8 | CAR 18:3 | MolDiscovery |
| 426.357_14.3 | CAR 18:1 | MassQL |
| 428.373_16.6 | CAR 18:0 | GNPS Library Match |
| 442.388_16.9 | CAR 19:0 | MS2LDA |
| 446.327_12.7 | CAR 20:5 | MassQL |
| 456.404_16.8 | CAR 20:0 | MS2LDA |
| 476.373_14.4 | CAR 22:4 | MolDiscovery |
| 484.436_18.1 | CAR 22:0 | MS2LDA |
| 510.451_18.2 | CAR 24:1 | MS2LDA |
| 512.467_19.1 | CAR 24:0 | MS2LDA |
| 536.467_18.7 | CAR 26:2 | MS2LDA |
| 598.504_19.5 | CAR 10:0-(O-18:0) | MassQL |
| 626.534_20 | CAR 14:0-(O-16:0) | MS2LDA |
| 636.519_19.7 | CAR 18:0-(O-13:2) | MassQL |
| 640.55_19.9 | CAR 13:0-(O-18:0) | MassQL |
| 648.519_19.7 | CAR 16:3-(O-16:0) | MassQL |
| 650.534_20 | CAR 14:2-(O-18:0) | MassQL |
| 652.551_20.1 | CAR 14:1-(O-18:0) | MassQL |
| 654.567_19.7 | CAR 16:0-(O-16:0) | MassQL |
| 664.55_19.1 | CAR 15:2-(O-18:0) | MS2LDA |
| 666.566_20.0 | CAR 13:0-(O-20:1) | MassQL |

| | | |
|---|---|---|
| 668.581_19.7 | CAR 15:0-(O-18:0) | MS2LDA |
| 676.55_19.2 | CAR 18:3-(O-16:0) | MassQL |
| 702.566_19.2 | CAR 16:0-(O-20:4) | MS2LDA |
| 728.581_19.9 | CAR 20:4-(O-18:1) | MassQL |
| **DGCC** | | |
| 798.567_18.3 | DGCC(16:0_(18:3-3OH)) | MassQL |
| 750.587_20.3 | DGCC(16:0_18:3) | MassQL |
| 574.395_14 | Unknown DGCC[#] | Manual $MS^2$ Annotation |
| 674.52_16.1 | DGCC(16:0_(11:0-OH)) | Manual $MS^2$ Annotation |
| 766.545_18.6 | DGCC(16:0_(17:4-2OH)) | Manual $MS^2$ Annotation |
| 774.584_19.6 | DGCC(16:0_20:5) | GNPS Library Match |
| 490.373_13.4 | Lyso-DGCC(16:0) | GNPS Library Match |
| **Pigments** | | |
| 551.424_19.4 | Echinenone* | GNPS Library Search |
| 551.425_18.9 | Echinenone* | GNPS Library Search |
| 567.418_18.6 | Diatoxanthin | GNPS Library Search |
| 583.415_18.1 | Diadinoxanthin or Diadinochrome* | Literature Search[2] |
| 583.415_20.5 | Diadinoxanthin or Diadinochrome* | Literature Search[2] |
| 585.43_17 | Antheraxanthin | Literature Search[2] |
| 597.394_17.8 | Astaxanthin | GNPS Library Hit |
| 601.423_15.1 | Neoxanthin | Literature Search[2] |
| 613.352_17.7 | Pyrroxanthin | Literature Search[2] |
| 631.363_15.5 | Peridinin | GNPS Library Search |
| 659.430_16.3 | Fucoxanthin | Analytical Standard[3] |
| 583.416_18.4 | Diadinoxanthin or Diadinochrome* | Literature Search[2] |
| [1]Deutsch *et al.* 2021 (3) | [2]Wakahama *et al.* 2012 (4) | [3]Level 1 Annotation, all other reported annotations are Level 2 (5) |
| [#]Contains 16:0 FA tail | *isomeric compounds with identical $MS^2$ spectra but different retention times | |

**Table S3.** MASST results for features annotated as acylated acylcarnitines and *N*-acyl taurine, where "X" indicates that feature contained a match from the dataset (massIVE ID indicated in the rightmost column).

| **Putative Annotation** | ***m/z*_RT** | ***Organism/Sample Type*** *Montipora capitata* | *Porites* | *Sarcophyton* spp. | *Montastraea cavernosa* | *Lobophytum* spp. | Tunicata | Marine sponges, algae and tunicates | Coral | **MassIVE IDs (MSV0000…)** |
|---|---|---|---|---|---|---|---|---|---|---|
| **Car 14:0-(O-16:0)** | **626.534_20** | X | | | | | | | | 85925 |
| **Car 13:0-(O-18:0)** | **640.55_19.9** | | X | X | X | X | X | | | 87449, 80632, 80597, 84587, 87471, 84430 |
| **Car 14:2-(O-18:0)** | **650.534_20** | | | | | | X | | | 84143 |
| **Car 14:1-(O-18:0)** | **652.551_20.1** | | X | | | | | | X | 80597, 80632, 79808 |
| **Car 16:0-(O-16:0)** | **654.567_19.8** | | | | | | X | X | | 87449 |
| **Car 15:0-(O-18:0)** | **668.581_19.7** | | | | | | X | | | 87449 |
| **Car 18:3-(O-16:0)** | **676.55_19.2** | | | | | | X | | | 87449, 84143 |
| **Car 16:0-(O-20:4)** | **702.566_19.3** | | | | | | X | | | 84143 |
| | | *Bacillariophyta* | *Lingulodinium* | *Pseudonitzschia* | *Synechococcus* | Coral Reef Seawater | Dissolved Organic Matter and Algal Extracts | | | |
| **Tau-C10:0** | **280.157_7.9** | X | X | X | X | | X | | | 85555, 87692, 81731, 88187, 87702, 87935, 88062, 87995, 88008, 87608, 87322, 87861, 86926, 85779, 85843, 83632, 83889, 85480, 86236, 87863, 88823, 85852, 88734, 88723, 82082, 82952, 86993, 87747 |

**Table S4.** Predicted chemical classes of features uniquely detected in BuOH partitions.

| ***m/z*_RT** | **Most Specific Class (CANOPUS)** |
|---|---|
| 230.248_12.4 | 1,2-aminoalcohols |
| 258.279_14.3 | 1,2-aminoalcohols |
| 286.31_17.3 | 1,2-aminoalcohols |
| 428.445_20.6 | 1,2-aminoalcohols |
| 523.472_17.4 | 1,2-diacylglycerols |
| 353.232_5.7 | 1,2-diacylglycerols |
| 304.176_4.8 | 1,3-substituted cyclopentyl purine nucleosides |
| 512.278_11.7 | 1-acyl-sn-glycero-3-phosphocholines |
| 492.272_10.1 | 1-acyl-sn-glycero-3-phosphocholines |
| 508.304_12 | 1-acyl-sn-glycero-3-phosphocholines |
| 496.339_23.9 | 1-acyl-sn-glycero-3-phosphocholines |
| 511.257_17.5 | 1-acyl-sn-glycero-3-phosphocholines |
| 564.365_11.2 | 1-acyl-sn-glycero-3-phosphocholines |
| 468.307_9.6 | 1-acyl-sn-glycero-3-phosphocholines |
| 552.401_11.9 | 1-acyl-sn-glycero-3-phosphocholines |

| | |
|---|---|
| 550.349_10.9 | 1-acyl-sn-glycero-3-phosphocholines |
| 542.323_10.5 | 1-acyl-sn-glycero-3-phosphocholines |
| 468.309_9.9 | 1-acyl-sn-glycero-3-phosphocholines |
| 572.334_10.9 | 1-acyl-sn-glycero-3-phosphocholines |
| 542.324_10.7 | 1-acyl-sn-glycero-3-phosphocholines |
| 742.574_18.4 | 1-alkyl,2-acylglycero-3-phosphocholines |
| 738.543_23.8 | 1-alkyl,2-acylglycero-3-phosphocholines |
| 524.37_23.9 | 1-alkyl,2-acylglycero-3-phosphocholines |
| 564.402_16 | 1-alkyl,2-acylglycero-3-phosphocholines |
| 812.585_20 | 1-alkyl,2-acylglycero-3-phosphocholines |
| 742.575_20.1 | 1-alkyl,2-acylglycero-3-phosphocholines |
| 469.342_17.9 | 3-alkylindoles |
| 344.279_12.4 | Acylcarnitines |
| 266.139_4.1 | Acylcarnitines |
| 484.436_20 | Acylcarnitines |
| 374.29_9.1 | Acylcarnitines |
| 422.326_13.7 | Acylcarnitines |
| 400.341_17 | Acylcarnitines |
| 428.372_18.1 | Acylcarnitines |
| 456.405_19.2 | Acylcarnitines |
| 476.374_17.3 | Acylcarnitines |
| 506.42_19.2 | Acylcarnitines |
| 504.404_18.6 | Acylcarnitines |
| 442.389_18.7 | Acylcarnitines |
| 428.372_18.4 | Acylcarnitines |
| 510.451_20.1 | Acylcarnitines |
| 484.435_20.1 | Acylcarnitines |
| 414.355_17.9 | Acylcarnitines |
| 398.326_16.1 | Acylcarnitines |
| 448.342_16 | Acylcarnitines |
| 193.195_14.8 | Alkatrienes |
| 295.172_14.6 | Alkyl glycosides |
| 357.213_7.3 | Alkyl glycosides |
| 760.536_19.6 | Alpha amino acid amides |
| 718.547_19.7 | Alpha amino acid amides |
| 324.167_5.7 | Alpha amino acid esters |
| 516.403_12.1 | Alpha amino acid esters |
| 504.389_16.1 | Alpha amino acids |
| 462.342_12.9 | Alpha amino acids |
| 520.346_10.9 | Alpha amino acids |
| 542.288_11.3 | Alpha amino acids |
| 504.39_17.5 | Alpha amino acids |
| 528.353_11.2 | Alpha amino acids |

| | |
|---|---|
| 514.373_13.8 | Alpha amino acids |
| 361.21_7.9 | Alpha amino acids |
| 472.363_17.8 | Alpha amino acids |
| 486.342_12.6 | Alpha amino acids |
| 722.555_21 | Alpha amino acids |
| 504.389_17.3 | Alpha amino acids |
| 472.363_17.8 | Alpha amino acids |
| 504.389_16.8 | Alpha amino acids |
| 414.356_17.6 | Alpha amino acids |
| 434.312_15.3 | Alpha amino acids |
| 412.377_17.7 | Alpha amino acids |
| 529.276_10.2 | Alpha amino acids and derivatives |
| 451.236_16.4 | Alpha amino acids and derivatives |
| 412.378_17.4 | Alpha amino acids and derivatives |
| 485.337_16 | Alpha amino acids and derivatives |
| 289.153_6 | Alpha amino acids and derivatives |
| 345.993_7 | Alpha amino acids and derivatives |
| 518.405_17.5 | Alpha amino acids and derivatives |
| 540.426_17.6 | Alpha amino acids and derivatives |
| 540.426_17.6 | Alpha amino acids and derivatives |
| 434.312_14.4 | Alpha amino acids and derivatives |
| 504.389_16.4 | Alpha amino acids and derivatives |
| 311.244_8 | Alpha amino acids and derivatives |
| 414.353_18.3 | Alpha amino acids and derivatives |
| 708.309_16.1 | Alpha amino acids and derivatives |
| 519.355_24 | Amino acids |
| 332.149_9 | Amino acids |
| 343.2_7.9 | Amino acids |
| 346.274_15.9 | Amino acids and derivatives |
| 282.059_9.5 | Amino acids and derivatives |
| 463.338_16 | Amino acids and derivatives |
| 285.29_11.5 | Amino acids and derivatives |
| 271.274_11.5 | Amino acids and derivatives |
| 313.321_12.8 | Amino acids and derivatives |
| 546.436_19 | Amino acids and derivatives |
| 369.384_18.1 | Amino acids and derivatives |
| 394.342_17.8 | Amino acids and derivatives |
| 402.323_17.1 | Amino acids and derivatives |
| 369.383_17.7 | Amino acids and derivatives |
| 500.394_18.5 | Amino acids and derivatives |
| 498.452_20.5 | Amino acids and derivatives |
| 309.178_7.1 | Amino acids and derivatives |
| 340.323_17.6 | Amino acids and derivatives |

| | |
|---|---|
| 426.394_20 | Amino acids and derivatives |
| 452.373_17.5 | Amino acids and derivatives |
| 418.317_16.9 | Amino acids and derivatives |
| 286.201_9.3 | Amino acids and derivatives |
| 294.157_6.5 | Amino acids, peptides, and analogues |
| 482.284_18.4 | Aminocyclitol glycosides |
| 368.153_19.4 | Aminotriazines |
| 296.237_17.5 | Aniline and substituted anilines |
| 248.143_9.1 | Aniline and substituted anilines |
| 527.217_11.1 | Anisoles |
| 246.128_8.6 | Aralkylamines |
| 307.161_6.8 | Aralkylamines |
| 380.354_19.6 | Azacyclic compounds |
| 165.091_20.3 | Benzaldehydes |
| 455.328_19.6 | Benzene and substituted derivatives |
| 345.278_16 | Benzene and substituted derivatives |
| 329.211_10 | Benzene and substituted derivatives |
| 340.214_8.7 | Benzene and substituted derivatives |
| 345.278_15.9 | Benzene and substituted derivatives |
| 340.248_7.1 | Benzene and substituted derivatives |
| 305.139_21.8 | Benzoic acids and derivatives |
| 327.122_5.3 | Benzoic acids and derivatives |
| 453.021_7.4 | Benzothiazoles |
| 339.936_5.7 | Bisphosphonates |
| 314.305_18.2 | Carboxylic acid amides |
| 480.384_19 | Carboxylic acid derivatives |
| 414.393_19.3 | Ceramides |
| 540.499_22.4 | Ceramides |
| 476.358_13.7 | Cholines |
| 506.404_14.3 | Cholines |
| 520.345_10.6 | Cholines |
| 506.404_14.1 | Cholines |
| 725.58_12.7 | Cyclic depsipeptides |
| 330.336_19.1 | Dialkyl ethers |
| 594.339_8.2 | Dicarboxylic acids and derivatives |
| 601.36_12.4 | Dicarboxylic acids and derivatives |
| 378.222_16.5 | Diphenylmethanes |
| 378.222_16.7 | Diphenylmethanes |
| 331.226_12.4 | Eicosanoids |
| 459.488_12.6 | Ethers |
| 259.19_12 | Fatty acid esters |
| 375.252_10.8 | Fatty acid esters |
| 447.332_15.8 | Fatty acid esters |

| | |
|---|---|
| 514.373_13.8 | Fatty acid esters |
| 444.404_17.3 | Fatty acid esters |
| 392.205_16.4 | Fatty acid esters |
| 512.467_21 | Fatty acid esters |
| 538.483_21 | Fatty acid esters |
| 530.368_12.3 | Fatty acid esters |
| 510.429_19.7 | Fatty acid esters |
| 521.492_20.4 | Fatty acid esters |
| 568.457_18.5 | Fatty acid esters |
| 339.23_12.1 | Fatty acid esters |
| 430.388_18.6 | Fatty acid esters |
| 390.284_8.2 | Fatty acids and conjugates |
| 547.331_19.7 | Fatty acyl glycosides |
| 836.602_20.6 | Fatty acyl glycosides of mono- and disaccharides |
| 407.352_20.1 | Fatty Acyls |
| 525.498_12.6 | Fatty Acyls |
| 400.378_18.5 | Fatty Acyls |
| 364.321_19.7 | Fatty Acyls |
| 398.363_19.1 | Fatty Acyls |
| 374.362_17.8 | Fatty Acyls |
| 365.231_7.3 | Fatty alcohols |
| 825.584_20.5 | Fatty alcohols |
| 588.512_15.9 | Fatty amides |
| 268.263_13.3 | Fatty amides |
| 314.305_17.6 | Fatty amides |
| 268.263_12.8 | Fatty amides |
| 426.373_20.7 | Fatty amides |
| 698.591_21.1 | Fatty amides |
| 784.557_18.8 | Glycerophosphocholines |
| 848.564_16.1 | Glycerophosphocholines |
| 800.588_23.9 | Glycerophosphocholines |
| 784.56_18.9 | Glycerophosphocholines |
| 838.576_19.3 | Glycerophosphocholines |
| 568.287_8.1 | Glycerophosphocholines |
| 654.427_17.9 | Glycerophosphocholines |
| 840.617_20.2 | Glycerophosphocholines |
| 612.386_10.7 | Glycerophosphocholines |
| 848.565_16.2 | Glycerophosphocholines |
| 824.649_12.8 | Glycerophosphocholines |
| 558.318_11.3 | Glycerophosphocholines |
| 772.572_21 | Glycerophosphocholines |
| 762.586_21.4 | Glycerophosphocholines |

| | |
|---|---|
| 574.314_10.2 | Glycerophosphocholines |
| 483.254_11.6 | Glycerophospholipids |
| 332.208_8.7 | Glycosylamines |
| 379.379_19.9 | Guanidines |
| 379.378_19.6 | Guanidines |
| 215.082_3.3 | Harmala alkaloids |
| 487.411_24.1 | Heteroaromatic compounds |
| 371.221_9.6 | Kaurane diterpenoids |
| 530.287_23.8 | L-alpha-amino acids |
| 260.222_11.5 | Leucine and derivatives |
| 507.294_10.2 | Lipids and lipid-like molecules |
| 540.427_16.8 | Lipids and lipid-like molecules |
| 318.226_7.3 | Long-chain fatty acids |
| 329.21_8.6 | Long-chain fatty acids |
| 528.272_11.2 | Lysophosphatidylcholines |
| 524.277_11.3 | Lysophosphatidylcholines |
| 558.319_12.1 | Lysophosphatidylcholines |
| 522.355_9.4 | Lysophosphatidylcholines |
| 506.287_10.2 | Lysophosphatidylethanolamines |
| 506.288_10.1 | Lysophosphatidylethanolamines |
| 588.446_15.6 | Macrolides and analogues |
| 286.201_9.4 | Methyl-branched fatty acids |
| 516.984_7.4 | Monoalkyl phosphates |
| 383.242_9.5 | Monosaccharides |
| 365.363_19.2 | Monoterpenoids |
| 329.316_13.3 | N-acyl amines |
| 301.285_11.6 | N-acyl amines |
| 240.232_11.6 | N-acyl amines |
| 546.436_18.8 | N-acyl amines |
| 574.467_20 | N-acyl amines |
| 329.316_12.9 | N-acyl amines |
| 329.316_12.8 | N-acyl amines |
| 396.31_13 | N-acyl amines |
| 518.405_17.8 | N-acyl amines |
| 500.394_18.8 | N-acyl amines |
| 468.441_19 | N-acyl amines |
| 440.409_17.6 | N-acyl amines |
| 568.457_18.9 | N-acyl amines |
| 440.409_16.9 | N-acyl amines |
| 708.577_23 | N-acyl amines |
| 468.441_19.1 | N-acyl amines |
| 574.467_19.9 | N-acyl amines |
| 383.373_17.2 | N-acyl amines |

| | |
|---|---|
| 380.217_9 | N-acyl amines |
| 554.55_20.4 | N-acyl amines |
| 301.286_12.2 | N-acyl amines |
| 440.409_17.7 | N-acyl amines |
| 398.363_19 | N-acyl-alpha amino acids |
| 346.258_8.2 | N-acyl-alpha amino acids |
| 344.242_8.8 | N-acyl-alpha amino acids |
| 300.216_6.9 | N-acyl-alpha amino acids |
| 546.4_17.6 | N-acyl-alpha amino acids |
| 532.421_18.2 | N-acyl-alpha amino acids |
| 604.406_16.1 | N-acyl-alpha amino acids |
| 571.431_14.8 | N-acyl-alpha amino acids |
| 546.399_17 | N-acyl-alpha amino acids |
| 344.242_9.3 | N-acyl-alpha amino acids |
| 358.258_9.6 | N-acyl-alpha amino acids and derivatives |
| 388.249_9.8 | N-acyl-alpha amino acids and derivatives |
| 345.203_5.9 | N-acyl-alpha amino acids and derivatives |
| 520.347_7.2 | N-acyl-alpha amino acids and derivatives |
| 570.34_7.7 | N-acyl-alpha amino acids and derivatives |
| 604.406_16.7 | N-acyl-alpha amino acids and derivatives |
| 518.405_18.3 | N-acyl-alpha amino acids and derivatives |
| 544.384_17.2 | N-acyl-L-alpha-amino acids |
| 544.384_16.9 | N-acyl-L-alpha-amino acids |
| 299.198_6.9 | N-acyl-L-alpha-amino acids |
| 590.391_16.3 | N-acyl-L-alpha-amino acids |
| 437.025_7 | Naphthalenes |
| 399.237_7.9 | Naphthalenes |
| 410.218_6.6 | N-carbamoyl-alpha amino acids and derivatives |
| 686.358_16.5 | Oligopeptides |
| 469.313_15.8 | Oligopeptides |
| 457.313_14.8 | Oligopeptides |
| 568.287_9.1 | Oligopeptides |
| 729.468_10.1 | Oligopeptides |
| 680.473_18.2 | Oligopeptides |
| 562.361_13.4 | Oligopeptides |
| 402.284_9.4 | Oligopeptides |
| 574.395_16.5 | Oligopeptides |
| 522.362_8.4 | Oligopeptides |
| 405.219_16.8 | Oligopeptides |
| 594.339_8.3 | Oligopeptides |
| 574.395_17 | Oligopeptides |
| 314.232_9.5 | Organic phosphonic acids |
| 326.199_8.1 | Organic phosphoramides |

| | |
|---|---|
| 438.503_21 | Organonitrogen compounds |
| 360.29_17.8 | Oxosteroids |
| 373.31_16.7 | Oxosteroids |
| 396.214_7.8 | Peptides |
| 574.358_10.2 | Peptides |
| 500.394_18.9 | Peptides |
| 616.442_17.4 | Peptides |
| 532.385_13.7 | Peptides |
| 496.333_9.4 | Peptides |
| 682.488_18.2 | Peptides |
| 506.404_16 | Peptides |
| 468.42_23 | Phenanthrenes and derivatives |
| 450.394_18 | Phenol ethers |
| 467.203_8.3 | Phenoxy compounds |
| 512.299_9.1 | Phenylalanine and derivatives |
| 618.421_17.4 | Phenylalanine and derivatives |
| 404.313_16.8 | Phenylalanine and derivatives |
| 388.393_19.1 | Phenylmethylamines |
| 360.362_18 | Phenylmethylamines |
| 332.331_17.1 | Phenylmethylamines |
| 332.331_15.7 | Phenylmethylamines |
| 332.331_16.4 | Phenylmethylamines |
| 360.362_18.4 | Phenylmethylamines |
| 416.425_20 | Phenylmethylamines |
| 846.606_20.1 | Phosphatidylcholines |
| 800.581_23.8 | Phosphatidylcholines |
| 826.594_23.8 | Phosphatidylcholines |
| 732.554_16.7 | Phosphatidylcholines |
| 732.554_17.2 | Phosphatidylcholines |
| 754.537_23.8 | Phosphatidylcholines |
| 828.609_23.8 | Phosphatidylcholines |
| 818.589_23.8 | Phosphatidylcholines |
| 764.573_23.9 | Phosphatidylcholines |
| 732.553_19.1 | Phosphatidylcholines |
| 732.553_19.1 | Phosphatidylcholines |
| 652.455_16.6 | Phosphatidylcholines |
| 688.497_17.9 | Phosphatidylcholines |
| 620.392_18.1 | Phosphatidylcholines |
| 786.601_17.2 | Phosphatidylcholines |
| 552.329_9.3 | Phosphatidylcholines |
| 580.397_23.9 | Phosphatidylcholines |
| 700.455_14.9 | Phosphatidylcholines |
| 612.386_11.4 | Phosphatidylcholines |

| | |
|---|---|
| 812.613_15.3 | Phosphatidylcholines |
| 732.554_18.6 | Phosphatidylcholines |
| 756.553_15.5 | Phosphatidylcholines |
| 756.553_15.8 | Phosphatidylcholines |
| 794.592_15.7 | Phosphatidylcholines |
| 806.568_14.5 | Phosphatidylcholines |
| 774.564_23.8 | Phosphatidylcholines |
| 554.344_8.8 | Phosphatidylcholines |
| 596.357_10.7 | Phosphatidylcholines |
| 580.397_15.4 | Phosphatidylcholines |
| 735.522_16.3 | Phosphatidylcholines |
| 808.584_20.8 | Phosphatidylcholines |
| 600.329_9 | Phosphatidylcholines |
| 822.564_19.8 | Phosphatidylcholines |
| 830.613_20.8 | Phosphatidylcholines |
| 846.617_18.3 | Phosphatidylcholines |
| 776.577_20.1 | Phosphatidylcholines |
| 800.578_21.1 | Phosphatidylcholines |
| 842.589_23.7 | Phosphatidylcholines |
| 582.34_12.8 | Phosphatidylethanolamines |
| 594.377_14 | Phosphatidylethanolamines |
| 702.473_16.4 | Phosphatidylethanolamines |
| 706.501_19.6 | Phosphatidylethanolamines |
| 528.272_10.3 | Phosphatidylethanolamines |
| 666.397_11.5 | Phosphatidylserines |
| 355.216_11.4 | Phosphinic acid esters |
| 542.344_9.7 | Phosphocholines |
| 531.366_16.6 | Phosphocholines |
| 610.334_7.3 | Phosphocholines |
| 674.519_18.6 | Phosphosphingolipids |
| 671.512_19.3 | Phosphosphingolipids |
| 701.559_23.9 | Phosphosphingolipids |
| 585.366_13 | Phosphosphingolipids |
| 456.275_8.5 | p-Hydroxybenzoic acid alkyl esters |
| 277.201_9.6 | Polyethylene glycols |
| 363.31_11.7 | Polyethylene glycols |
| 581.462_16.5 | Polyethylene glycols |
| 553.43_14.9 | Polyethylene glycols |
| 306.243_11.2 | Prenol lipids |
| 351.348_18.9 | Prenol lipids |
| 379.173_3 | Purine nucleosides |
| 423.01_6.7 | Pyrimidine nucleotides |
| 539.289_17.1 | Saccharolipids |

| | |
|---|---|
| 307.227_10.2 | Steroid acids |
| 464.374_19.7 | Steroid acids |
| 432.347_19.2 | Steroid esters |
| 424.341_16.2 | Steroidal alkaloids |
| 496.415_19.4 | Steroidal alkaloids |
| 456.42_22.9 | Steroidal alkaloids |
| 416.336_13.4 | Steroidal alkaloids |
| 373.31_13.6 | Steroids and steroid derivatives |
| 428.388_21.5 | Steroids and steroid derivatives |
| 442.404_22.1 | Steroids and steroid derivatives |
| 440.389_20.8 | Steroids and steroid derivatives |
| 246.128_8.6 | Styrenes |
| 403.209_10.2 | Styrenes |
| 767.497_17.2 | Sulfoquinovosyldiacylglycerols |
| 765.481_16.5 | Sulfoquinovosyldiacylglycerols |
| 274.159_14.3 | Triarylamines |
| 553.268_10.5 | Tricarboxylic acids and derivatives |
| 437.341_10.2 | Triterpenoids |
| 674.519_18.3 | Very long-chain fatty acids |
| 505.497_20.1 | Wax monoesters |
| 341.305_17.1 | - |
| 318.128_16.2 | - |
| 886.617_20.5 | - |
| 395.783_24.1 | - |
| 258.279_13.8 | - |
| 501.339_14.6 | - |
| 629.366_13.3 | - |
| 300.992_13.2 | - |
| 373.274_10.1 | - |
| 357.347_14.1 | - |
| 270.315_17.4 | - |
| 308.15_18.3 | - |
| 417.3_19.2 | - |
| 376.313_24 | - |
| 160.17_4.8 | - |
| 371.28_23.9 | - |
| 272.222_12.3 | - |
| 228.268_13.5 | - |
| 258.279_14 | - |
| 200.237_12.1 | - |
| 268.001_6.8 | - |
| 286.31_17 | - |
| 221.014_6.2 | - |

| | |
|---|---|
| 348.144_9 | - |
| 285.16_5.9 | - |
| 366.155_9 | - |
| 220.015_6.2 | - |
| 385.378_18.1 | - |
| 260.003_6.2 | - |
| 248.143_9.2 | - |
| 313.321_12.9 | - |
| 222.013_6.2 | - |
| 429.318_11.8 | - |
| 229.202_5.1 | - |
| 241.203_9.6 | - |
| 252.996_6 | - |
| 164.645_5.2 | - |
| 268.001_6.3 | - |
| 244.008_6.5 | - |
| 213.016_5.5 | - |
| 270.315_15.6 | - |
| 286.31_16.8 | - |
| 182.106_5.3 | - |
| 362.253_8 | - |
| 381.225_5.3 | - |
| 473.344_11.9 | - |
| 228.268_13.9 | - |
| 382.2_6.9 | - |
| 241.203_9.8 | - |
| 221.014_6.3 | - |
| 242.284_14 | - |
| 395.783_24.1 | - |
| 246.243_9.4 | - |
| 427.389_14.8 | - |
| 598.263_8.4 | - |
| 355.262_10.1 | - |
| 815.482_16.1 | - |
| 402.386_19.8 | - |
| 270.315_17 | - |
| 499.396_17.9 | - |
| 541.277_12.3 | - |
| 575.503_23.8 | - |
| 213.016_5.3 | - |
| 206.009_5.2 | - |
| 212.017_5.2 | - |
| 242.248_16.8 | - |

| | |
|---|---|
| 260.003_5.9 | - |
| 205.665_7.7 | - |
| 164.644_5 | - |
| 244.008_6.1 | - |
| 374.291_13.8 | - |
| 252.996_5.8 | - |
| 313.273_23.8 | - |
| 480.331_11.7 | - |
| 319.263_15.5 | - |
| 383.79_23.9 | - |
| 271.263_24.9 | - |
| 212.017_5.6 | - |
| 214.015_5.7 | - |
| 236.01_6 | - |
| 221.007_5.2 | - |
| 206.009_5.5 | - |
| 245.007_6.5 | - |
| 206.037_5.1 | - |
| 191.003_4.6 | - |
| 214.253_12.2 | - |
| 241.203_10.1 | - |
| 176.036_4.5 | - |
| 258.279_16.2 | - |
| 206.037_5.4 | - |
| 157.092_6 | - |
| 281.009_4.8 | - |
| 498.342_9.3 | - |
| 286.31_17.9 | - |
| 314.232_9.5 | - |
| 344.352_20.5 | - |
| 366.265_11.4 | - |
| 316.248_10.4 | - |
| 267.027_10.3 | - |
| 353.744_15.1 | - |
| 280.3_17.7 | - |
| 411.769_24.2 | - |
| 250.642_10.4 | - |
| 371.722_15.8 | - |
| 268.001_6.6 | - |
| 298.346_18.9 | - |
| 201.16_4.7 | - |
| 314.341_19 | - |
| 360.237_7.1 | - |

| | |
|---|---|
| 200.237_12 | - |
| 380.211_7.3 | - |
| 404.786_24.1 | - |
| 260.258_10.4 | - |
| 304.284_12.4 | - |
| 268.671_10.9 | - |
| 333.349_21.4 | - |
| 292.192_24 | - |
| 341.187_8.6 | - |
| 999.756_14.6 | - |
| 289.153_5.2 | - |
| 239.225_12.2 | - |
| 320.699_24 | - |
| 425.287_24.2 | - |
| 263.684_24.1 | - |
| 389.254_24.1 | - |
| 291.302_19.2 | - |
| 484.762_5.9 | - |
| 268.176_24.1 | - |
| 354.844_24.3 | - |
| 268.263_13 | - |
| 526.483_21.2 | - |
| 396.456_21.5 | - |
| 206.009_5.9 | - |
| 244.999_5.9 | - |
| 765.749_7.1 | - |
| 342.278_23.9 | - |
| 209.14_4.1 | - |
| 414.191_7.1 | - |
| 382.199_6.7 | - |
| 372.383_23.3 | - |
| 372.383_23.4 | - |
| 371.722_15.2 | - |
| 244.21_24 | - |
| 929.577_21 | - |
| 1593.361_10.8 | - |
| 1199.27_10.8 | - |
| 389.321_23.7 | - |
| 332.165_3.6 | - |
| 1191.271_11 | - |
| 252.207_24.1 | - |
| 258.279_15.8 | - |
| 420.757_24.1 | - |

| | |
|---|---|
| 238.191_24.1 | - |
| 296.258_17.4 | - |
| 1199.269_10.7 | - |
| 323.158_4.5 | - |
| 248.193_7.3 | - |
| 426.393_20.6 | - |
| 433.763_24.1 | - |
| 850.594_23.8 | - |
| 584.282_6.4 | - |
| 382.331_23.9 | - |
| 383.79_20.3 | - |
| 244.21_21.4 | - |
| 286.31_17.5 | - |
| 239.225_16.7 | - |
| 296.684_24.1 | - |
| 858.582_15.6 | - |
| 422.399_22.5 | - |
| 852.608_23.8 | - |
| 389.321_22.6 | - |
| 358.367_21.9 | - |
| 358.368_22.6 | - |
| 1109.723_9.9 | - |
| 536.755_6.5 | - |
| 255.182_6.1 | - |
| 441.198_7.4 | - |
| 403.336_24 | - |
| 1082.454_10.6 | - |
| 860.6_23.8 | - |
| 454.404_22.2 | - |
| 402.328_22.6 | - |

**Table S5.** Symbiodiniaceae strains cultured in this study (strains were originally sourced from the BURR collection and kindly provided by Mary Alice Coffroth).

| Genera | Strain ID from BURR Collection |
|---|---|
| *Breviolum* | *Breviolum* MF 1.05b.01 SCI 07-205 |
| *Breviolum* | *Breviolum* MF 1.05b.01 SCI 07-209 |
| *Breviolum* | *Breviolum* Mf 10.14.02 SCI |
| *Durusdinium* | *Durusdinium* Mf 2.2b-2 |
| *Durusdinium* | *Durusdinium* Mf 2.2b-2 |
| *Symbiodinium* | *Symbiodinium* Mf 10.02a |
| *Symbiodinium* | *Symbiodinium* 04-503 SCI.01 |
